# Non-photosynthetic Plastid Replacement by a Primary Plastid in the Making

**DOI:** 10.1101/2025.10.07.680570

**Authors:** Yong Heng Phua, Dewi Langlet, Bruno M. Humbel, Arno Hagenbeek, Vasyl Vaskivskyi, Kevin Wakeman, Filip Husnik

## Abstract

The integration of a cyanobacterium into a heterotrophic eukaryote gave rise to the primary plastid ∼1.5 Gya. This rare cyanobacterium-to-plastid transition has only been reported once more in *Paulinella chromatophora* ∼100 Mya. Unfortunately, the order and relative importance of organellogenesis events have been blurred by time in primary plastids and obscured by *P. chromatophora* becoming phototrophic. Here, we characterize the tripartite symbiosis in a benthic dinoflagellate (*Sinophysis* sp.) using diverse single-cell methods. *Sinophysis* houses a photosynthetic cyanobacterium closely interacting with an alphaproteobacterial endosymbiont. The cyanobacterium is in the intermediate stage of symbiont-to-organelle transition, with the host likely supporting it with metabolites and proteins, and controlling its cell division. Surprisingly, it seems to have replaced the host’s remnant non-photosynthetic plastid. Our results support mixotrophy, horizontal gene transfer, co-symbioses, and cell division control as early drivers of primary plastid origin and highlight the importance of protists for deciphering organellogenesis events.

## INTRODUCTION

Endosymbiosis between microorganisms has triggered the origin of novel lineages throughout the history of life. Two prominent examples are the endosymbiont-derived organelles, mitochondria and plastids, which have become highly integrated into eukaryotic cells^1^. Since the free-living ancestors of mitochondria and plastids were acquired approximately 2 billion years ago (Gya) and 1.5 Gya^2^ respectively, reconstructing major organellogenesis steps is limited by the lack of intermediate stages. Key events that frame this process are metabolite support by the host, horizontal gene transfer to the host genome from diverse donors (HGT) or the current endosymbiont (EGT), symbiont genome reduction, host control of the symbiont division, and protein targeting establishment^3^’^11^.

Although secondary, tertiary, and other complex plastid symbioses are common across eukaryotes, the uptake and subsequent establishment of a cyanobacterium into a primary plastid is comparatively rare^12^. Apart from the ancestor of Archaeplastida (green algae, red algae, and glaucophytes), the only such newly recorded plastid acquisition is the chromatophore established in the photosynthetic clade of the rhizarian amoeba *Paulinella* ∼100 Mya^13^. The chromatophore is highly integrated and displays key organelle features, such as genome reduction, protein targeting, and synchronised cell division with the host cell^4,9,14^ Interestingly, most gene transfers on the *Paulinella chromatophora* genome are the result of HGT from diverse bacterial donors rather than from the present day symbionts. *P. chromatophora* is fully phototrophic, but its heterotrophic relatives that do not contain chromatophores are bacterivorous. It is hypothesised that the ancestor of the photosynthetic *Paulinella* clade was mixotrophic for tens of millions of years, allowing for the continuous exposure to foreign DNA and eventual HGTs supporting the gene loss from the chromatophore genome and its integration^15^. Such a mixotrophic lifestyle is also seen in other protist hosts with complex plastids, such as dinoflagellates^16^.

Dinoflagellates are not only extremely versatile in acquiring plastids and photosymbionts from diverse donors, but there have also been multiple plastid losses and secondary/tertiary plastid replacements reported over the course of dinoflagellate evolution^17^. Additionally, many groups remain mixotrophic regardless of the presence of a photosynthetic plastid^16^. The family Dinophysiaceae comprises multiple genera that have lost their secondary photosynthetic plastid, but still retain a non-photosynthetic remnant of the organelle^18,19^ Some of them have acquired alternative ways to capture sunlight energy, such as kleptoplasty^20^ or ‘gardening’ cyanobacterial ectosymbionts in specialized extracellular chambers^21^. The benthic dinoflagellates from the genus *Sinophysis* have similarly lost their ancient photosynthetic plastids, and their ancestor was likely heterotrophic^22^. Interestingly, some species harbour intracellular green-brown bodies previously identified as cyanobacteria using 16S rRNA gene phylogenies^23^. However, the role of the cyanobacteria and the extent to which they are integrated with the dinoflagellate hosts have not been thoroughly examined.

Here, we show that *Sinophysis* represents a promising model of the initial stage of primary plastid integration. We confirm that it houses two endosymbionts, a cyanobacterium and an alphaproteobacterium, and report the extent of their genome reduction based on single-cell genomics. Following the identification of genes related to photosynthesis in the cyanobacterial genome, the photosynthetic ability of the host-symbiont system was confirmed with nanorespirometry. Volume electron microscopy identified physical interactions between the symbionts and the host, and their shared localization to a host-derived compartment (symbiosome) surrounded by the host mitochondria. Finally, single-cell transcriptomics revealed potential gene transfers on the dinoflagellate genome, a limited presence of plastid-related genes, and adaptations for long-term symbiont maintenance. Our results suggest that the cyanobacterial symbionts/chromatophores of *Sinophysis* are in the early-to intermediate stage of becoming primary plastids.

## RESULTS

### Localization and identification of the dual intracellular symbionts

Endosymbiotic cyanobacteria can be observed as multiple brown-green bodies within each *Sinophysis* sp. dinoflagellate cell (Figure 1a, b). The identity of the endosymbionts was confirmed with 16S rRNA-specific probes designed from the genome data generated in this study (Figure 1a, c, and **S1**). With electron microscopy, the cyanobacteria were observed in the periphery of the dinoflagellate cell and contained an internal thylakoid-like membrane (Figure 1d, e). Additionally, another bacterial endosymbiont is often found closely associated with the cyanobacterium in the same host-derived membrane-bound compartment resembling a symbiosome (Figure 1f). These bacteria were identified as Alphaproteobacteria from the *Nisaea* genus (Rhodospirillales) based on their 16S rRNA gene sequences being present in all 18 of our single-cell transcriptomes from two lineages of *Sinophysis* (Figure S2, S3, S4, Table S1).

**Figure 1.**
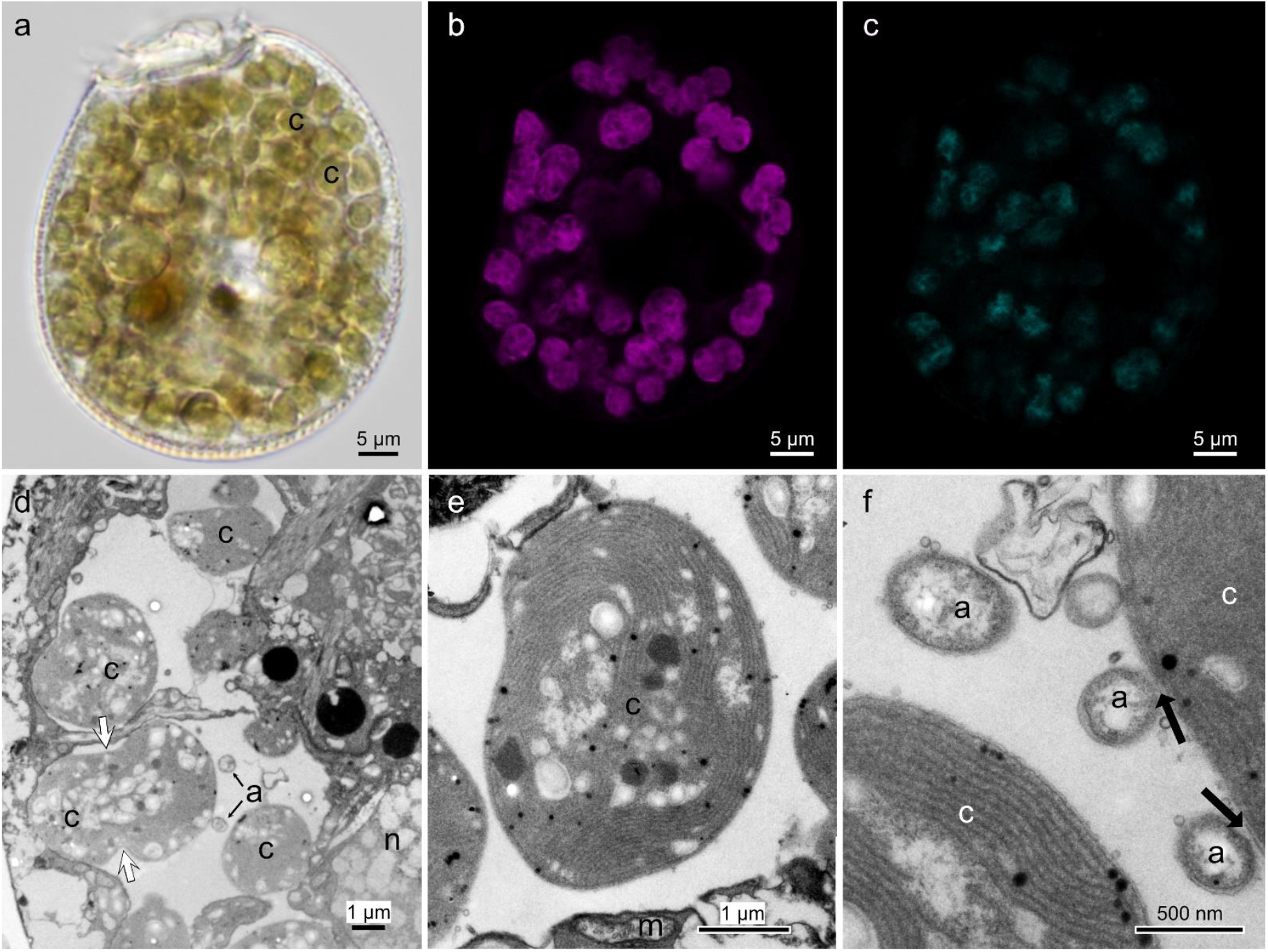
Micrographs of *Sinophysis* sp. **a)** Light micrograph. **b)** Laser scanning confocal micrograph of the autofluorescence from cyanobacterial symbionts (purple, excitation λ: 553 nm). **c)** Cell stained with cyanobacterial symbiont-specific 16S rRNA probe (cyan, excitation λ: 353 nm). **d)** TEM section showing the localization of cyanobacterial (c) and alphaproteobacterial symbionts (a), nucleus (n). **e)** High-magnification TEM image of the cyanobacterial symbiont showing thylakoid-like structures and dinoflagellate mitochondria (m) in close vicinity. **f)**. High-magnification image of the alphaproteobacterial symbionts in contact with cyanobacteria (black arrows).

### Genome features of the cyanobacterial symbiont

Total DNA from pooled cells isolated from field samples was sequenced, and genome statistics of both symbionts are shown in Table S2. The cyanobacterial genome consists of the main chromosome and two plasmids, totalling 2,178,118 bp with a 34.7% GC content (Figure 2, Table S2). The genome size is approximately half of the closest free-living relative *(Geminocystis herdmanii*, NCBI accession ID: NZ_CM00l 775.1, 4,263,418 bp). It contains 1,678 intact genes, 199 predicted pseudogenes, 42 tRNAs, 3 rRNA operons (3 sets of 23S and 16S rRNA genes), and 13 other non-coding RNAs. Genes related to photosynthesis are distributed throughout the main chromosome (Figure 2a) and the larger plasmid (Figure 2b). The smaller plasmid contained only a cyanobacterial chaperone gene (*dna*.*J*), helicase (dbpA), three viral genes with DNA-interacting domains, and 14 hypothetical proteins (Figure 2c). Analysis of the 16S rRNA gene places the cyanobacterial symbiont in the order Chroococcales, while diverse cyanobacteria symbiotic with dinoflagellates from the family Dinophysaceae originated from the genus *Synechococcus* within the order Synechococcales (Figure S5).

**Figure 2.**
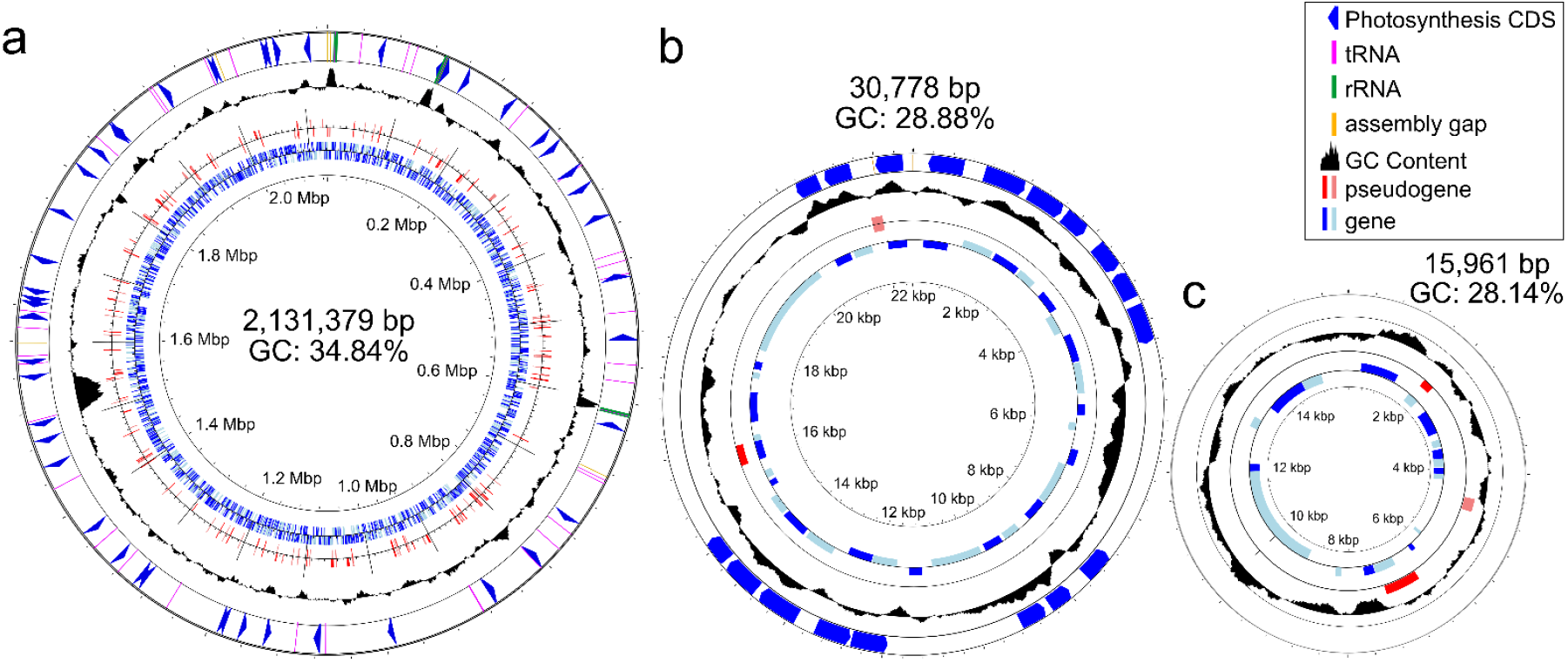
Graphical representation of the *Geminocystis photosymbiotica* genome. **a)** Cyanobacterial genome map. **b)** larger plasmid, **c)** smaller plasmid. From the outer circle to the inner circle depict: photosynthesis genes (blue arrow block), GC content, pseudogenes (red block), protein-coding genes (blue block).

To get an overview of the gene functions retained, the genes were divided into 25 Clusters of Orthologous Genes (COG) categories. Gene losses were disproportional compared to G. *herdmanii* (Figure S6a, b). The categories related to cell motility [N], signal transduction mechanisms [T], cell cycle/division [D], defence mechanisms [V], and replication, recombination and repair [L] had the highest percentage of gene loss(>70%). Core cellular functions such as glycolysis, carbon fixation, pyruvate oxidation, amino acid synthesis, and aminoacyl-tRNA synthesis are intact (Figure 3a). However, several hallmark gene losses indicative of an endosymbiotic lifestyle have occurred: *uvrABC* nucleotide excision repair genes involved in DNA repair have been lost, and the *thyX* pyrimidine deoxyribonucleotide biosynthesis gene is pseudogenised. In addition, genes related to the TCA cycle, succinic semialdehyde dehydrogenase *(gabD)*, and malate dehydrogenase (*mdh*)were not found in the genome. Biosynthesis of some cell envelope components, such as peptidoglycan (*mrcA. mrdA, vanX. rlpA*), lipopolysaccharides. ADP-Lglycero-Dmannoheptose, S-layer glycoprotein, and Type I bacterial secretion system (*hlyBD*). has also been reduced. Additionally, genes related to nitrogen metabolism have also been lost; the assimilatory nitrate reduction pathway (*nrt*ABCD) is entirely absent, and genes responsible for the biosynthesis of molybdenum cofactor are also lost when compared to the closest free-living relative. Since the closest free-living relatives are not nitrogen fixers, nitrogenase genes were likely never present rather than being lost.

**Figure 3.**
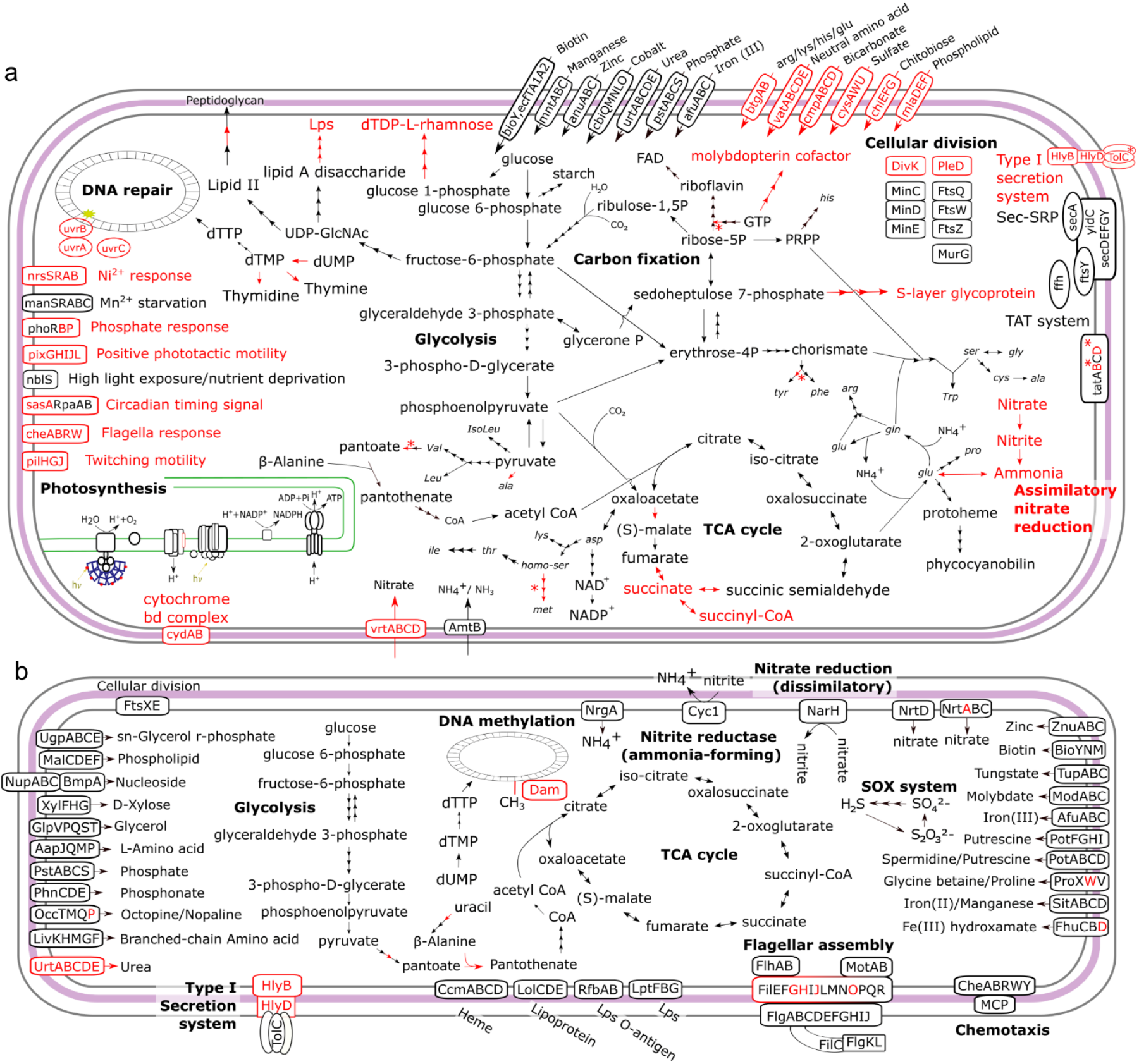
Cellular overviews with selected metabolic pathways of the two *Sinophysis* sp. endosymbionts as reconstructed from their genomes. **a)** Schematic of pathways present in the cyanobacterial symbiont. **b)** Schematic of pathways present in the alphaproteobacterial symbiont. Triple arrows indicate multiple reactions, and red arrows and text indicate enzymes, compounds, or pathways missing from the endosymbiont genomes but present in the free-living sister lineage *Geminocystis herdmanii* PCC 6308 and *Nisaea acidiphila*, respectively. Asterisks (*) indicate enzymes present neither in the endosymbiont nor in its free-living relatives.

### Genome features of the alphaproteobacterial symbiont

A draft genome of the alphaproteobacterial symbiont was assembled and binned into 211 contigs, totalling 4,435,508 bp with 64.0% GC content and CheckM scores of completeness 99.12, contamination 0.44, and strain heterogeneity 0.00. The binned genome consists of 3,613 intact genes and 996 pseudogenes when using *Nisaea acidiphila*, the closest available free-living relative, for pseudogene prediction (Figure S3). All COG categories contain pseudogenes, but genes in various transport/metabolism [Q, E, P, I], cellular signalling [N, T, V] categories, and transcription [K] showed high rates (>75%) of pseudogenisation (Figure S6c, d). Additionally, DNA adenine methylase *(dam)* was not found. DNA polymerase *dnaE2*, recombination protein *(recT)*, paralogs of exodeoxyribonuclease III *(xthA)*, and single-stranded DNA-binding proteins *(ssb)* have likely been pseudogenised. When compared with *N. acidiphila*, most core metabolic pathways, including DNA replication, glycolysis, pentose phosphate pathway, and amino acid biosynthesis, were retained. Additionally, some genes for flagella assembly have been lost. Receptors for chemotaxis and many transporters are present in the genome. Interestingly, the symbiont has also retained pathways that enable an anaerobic lifestyle, nitrate reduction, and pyruvate fermentation (Figure 3b).

### Maintenance of photosynthetic function despite drastic genome reduction

Analysis of the cyanobacterial genome revealed that most genes related to photosynthesis have been retained. Genes encoding photosystem I and II, photosynthetic electron transport, cytochrome b6/f complex, F-type ATPase, cyanobacterial antenna proteins (allophycocyanin, phycocyanin, and phycoerythrin), and haem biosynthesis were all present. From these proteins, PetL (Cytochrome b6-f complex subunit 6) and phycocyanin-associated linker protein CpcD were not identified. Due to the presence of photosynthesis-related genes (Figure 3a) and carboxysomes (Figure 1e) in the cyanobacterium, we tested the capability of the host-symbiont system to photosynthesize by examining the production of oxygen in the presence of light. Four batches of *Sinophysis* cells were evaluated for their photosynthetic capability at different light intensities. On average, the net photosynthesis rates were −21.2, 108.1, and 106.2 pmol*O*_2_/cell/day at 0, 115, and 300 μmol/m^2^/s light intensities, respectively, indicating that the cells were photosynthetically active (Figure S7). Linear models showed a significant effect of light intensity on the photosynthesis rate (df = 2,9; F = 15.9; p = 0.001) such as the photosynthesis rate is significantly lower in the dark than in the light (t = - 4.9; p < 0.001) and that there is no significant difference between 115 and 300 μmol/m^2^/s (t= 0.07, p = 0.94). Photosynthetic rates of *Prorocentrum* used as a control were in the same order of magnitude (− 26.7 pmol02/cell/day in the dark, 54.7 and 59.7 pmol*O*_2_/cell/day at 115 and 300 *µ*mol*photons*/m^2^/s, respectively).

### Tight association between two endosymbionts and

#### Sinophysis

The cyanobacteria were spherical or joined together in a dumbbell shape, forming clusters along the periphery of the host cell (Figure 4a, Movie Sl, S2). The alphaproteobacterial symbiont is located in the same symbiosome and in close contact with the cyanobacteria (Figure S8). Most symbionts appear to co-occur in a single symbiosome (Figure S9), but our data do not allow us to rule out the presence of multiple symbiosomes. Some cyanobacteria have a different electron density, and their thylakoid structures appear to be damaged (Figure S9b). Interestingly, all the cyanobacteria were in contact with the alphaproteobacterial symbiont (Figure 4a, b, c), but not all alphaproteobacteria were associated with cyanobacteria.

**Figure 4.**
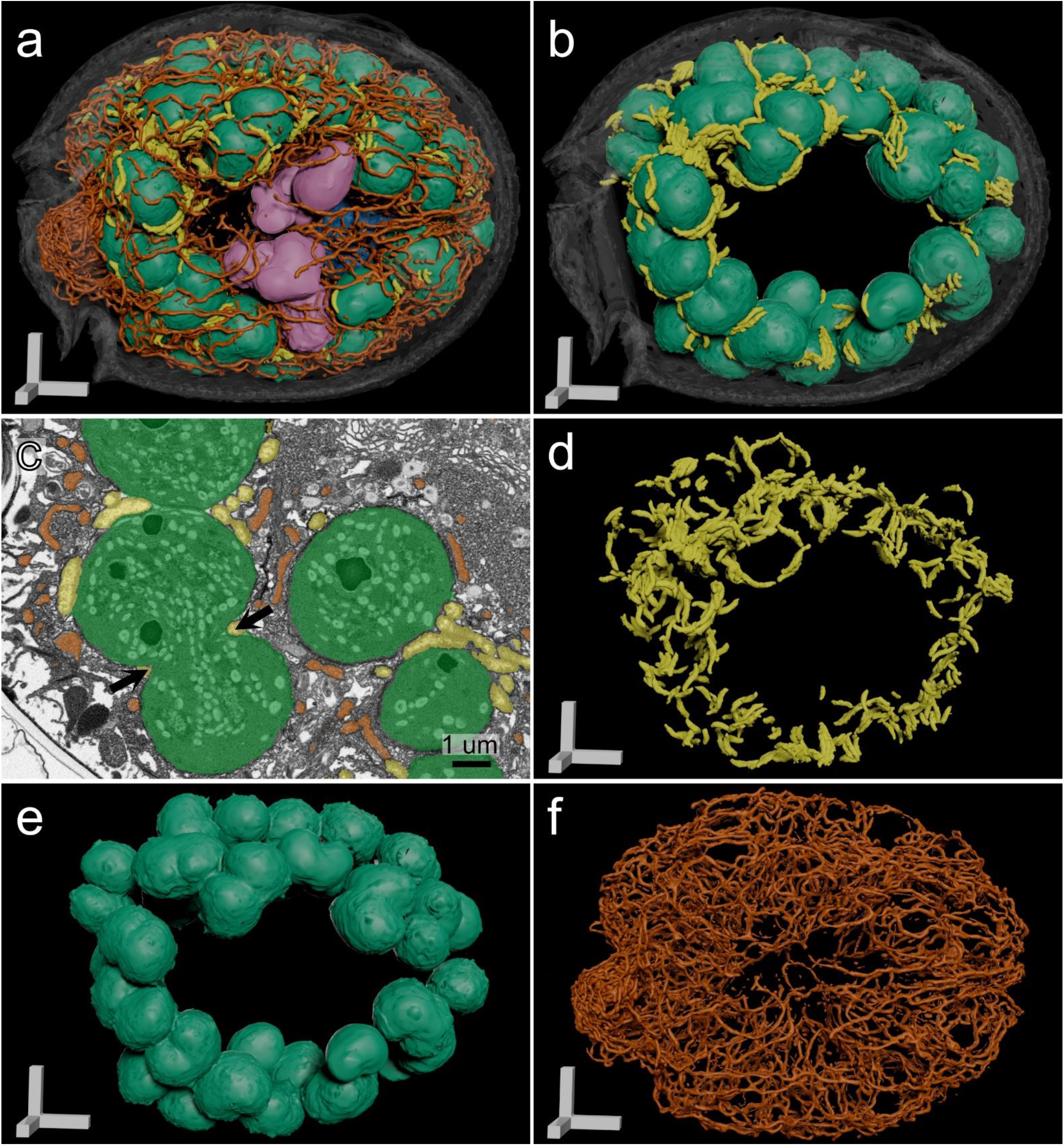
Three-dimensional reconstruction of Sinophysis sp. using focused ion beam scanning electron microscopy (FIB-SEM). **a)** Reconstruction of a whole cell showing the cyanobacteria (green), alphaproteobacteria (yellow), food vacuoles (purple), nucleus (blue), and mitochondria (orange). **b)** Segmentation showing the interactions of the cyanobacteria (green) and alphaproteobacteria (yellow). **c)** Slice of the cell highlighting the close association between the two symbionts, cleavage furrow (black arrow). Segmentation of **d)** alphaproteobacteria, **e)** cyanobacteria, and **f)** mitochondria. Scale bar = 5 µm unless stated otherwise (c).

Outside of the symbiosome, mitochondria form a reticular network in close proximity to the symbiosome (Figure 4a, c, f, and S9d). The mitochondrial network also forms a dense cluster around the epitheca (Figure 4a, f, likely due to the ATP requirements of the dinoflagellate flagella. Several food vacuoles are localised within the centre of the cell. The food vacuoles have electron-dense membranes that are distinct from the cyanobacterial symbionts, as well as clusters of minute crystalline structures (Figure S10). Volumetric analyses show that the symbionts represent almost a quarter of the total cell volume, comprising 23.3% cyanobacteria and 1.3% alphaproteobacteria in one fully imaged cell. In a second cell, where only a small portion of the host is imaged, the cyanobacteria are 30% of the total cell volume (Figure S11).

#### Absence of non-photosynthetic plastid-targeted genes from the dinoflagellate transcriptome

Two sets of single-cell transcriptomes, consisting of 4 and 5 cells, were co-assembled based on their near-identical 18S rRNA gene sequences. The BUSCO score (against alveolata_odblO) for sp. 1 was 123 complete (71.9%, [Single copy: 45.0%, Duplicated: 26.9%]) with 15 fragmented (8.8%) and 33 missing (19.3%) genes (n: 171), while sp. 2 had 78 complete (45.6% [Single-copy: 18.7%, Duplicated: 26.9%]), with 51 fragmented (29.8%) and 42 missing (24.6%) genes (n: 171). Most genes that are typically retained in non-photosynthetic plastids (biosynthesis of isoprenoids, haem, and fatty acids, Fe-S cluster assembly, and genes encoding the Translocon on the Inner Chloroplast membrane complex TIC) were not expressed in *Sinophysis* (5 out of 35 detected; Figure 5). However, other dinoflagellates retain most of these genes regardless of lifestyle. Interestingly, homologues of most of these genes are present in the genome of the cyanobacterial symbiont. In contrast, genes for mitochondria-targeted proteins were found to be retained similarly to other dinoflagellate relatives (Figure S12).

**Figure 5.**
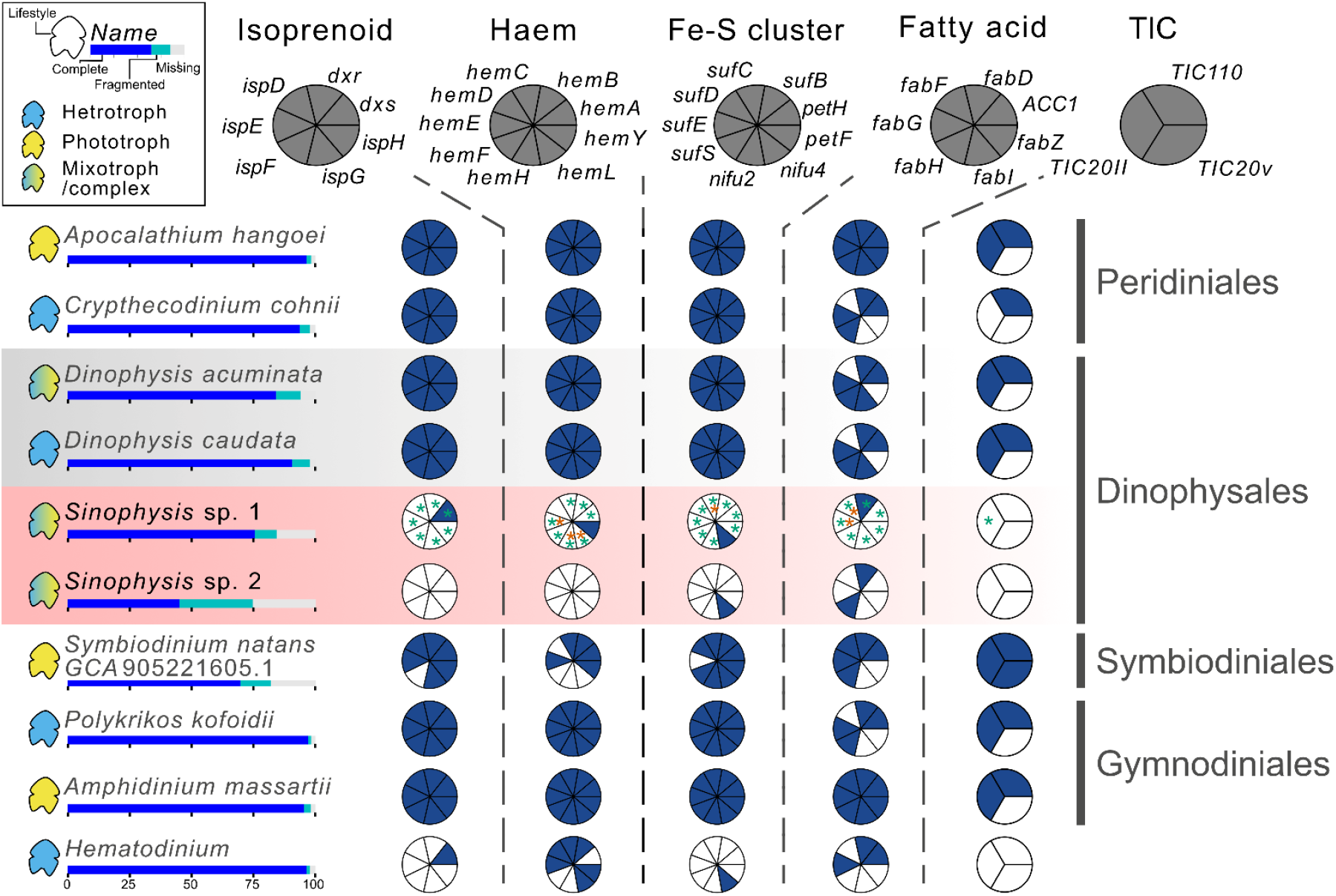
Transcriptome-based analysis of plastid-targeted proteins. Schematic depiction of several biosynthetic pathways commonly retained in non-photosynthetic plastids. Blue segments represent dinoflagellate genes that are present, white segments represent absent genes, and asterisks (*) represent homologues found in the symbionts (green asterisks represent cyanobacterial genes, and orange asterisks represent alphaproteobacterial genes).

As the ancestors of *Sinophysis* contained peridinin plastids of red algal secondary origin, we tested whether organelle-targeting mechanisms could have been adapted for the new symbiont. However, only a few genes were predicted to be targeted to hypothetical plastids, and there was no consensus between the three algorithms we used (Figure S13a). On the other hand, the same algorithms were able to identify plastid-targeted proteins from *Prorocentrum* minimum HBJL01 (Figure S13b) and *Dinophysis acuminata* GKBP01 (Figure S13c). Mitochondria-targeted proteins were also identified (Figure S13), confirming that there was no bias against organelle-targeting proteins in our data.

#### The dinoflagellate transcriptome contains genes acquired horizontally from diverse origins

To identify potential gene transfer candidates to *Sinophysis*, we screened the *de novo* assembled reference transcriptomes for proteins of bacterial origin. From the high-confidence transcripts, proteins present in the symbiont genomes were filtered out, resulting in a total of 313 genes identified (Figure S14, Table S3). Out of these, 165 genes were classified into phyla and lower taxonomic levels (Figure 6a). These genes originate from a wide variety of bacterial groups. However, the largest contributing group was Proteobacteria, specifically Alpha- and Gammaproteobacteria (Figure 6b, c). The functions of the HGTs were mostly linked to transport and metabolism (Figure S14, Table S3).

**Figure 6.**
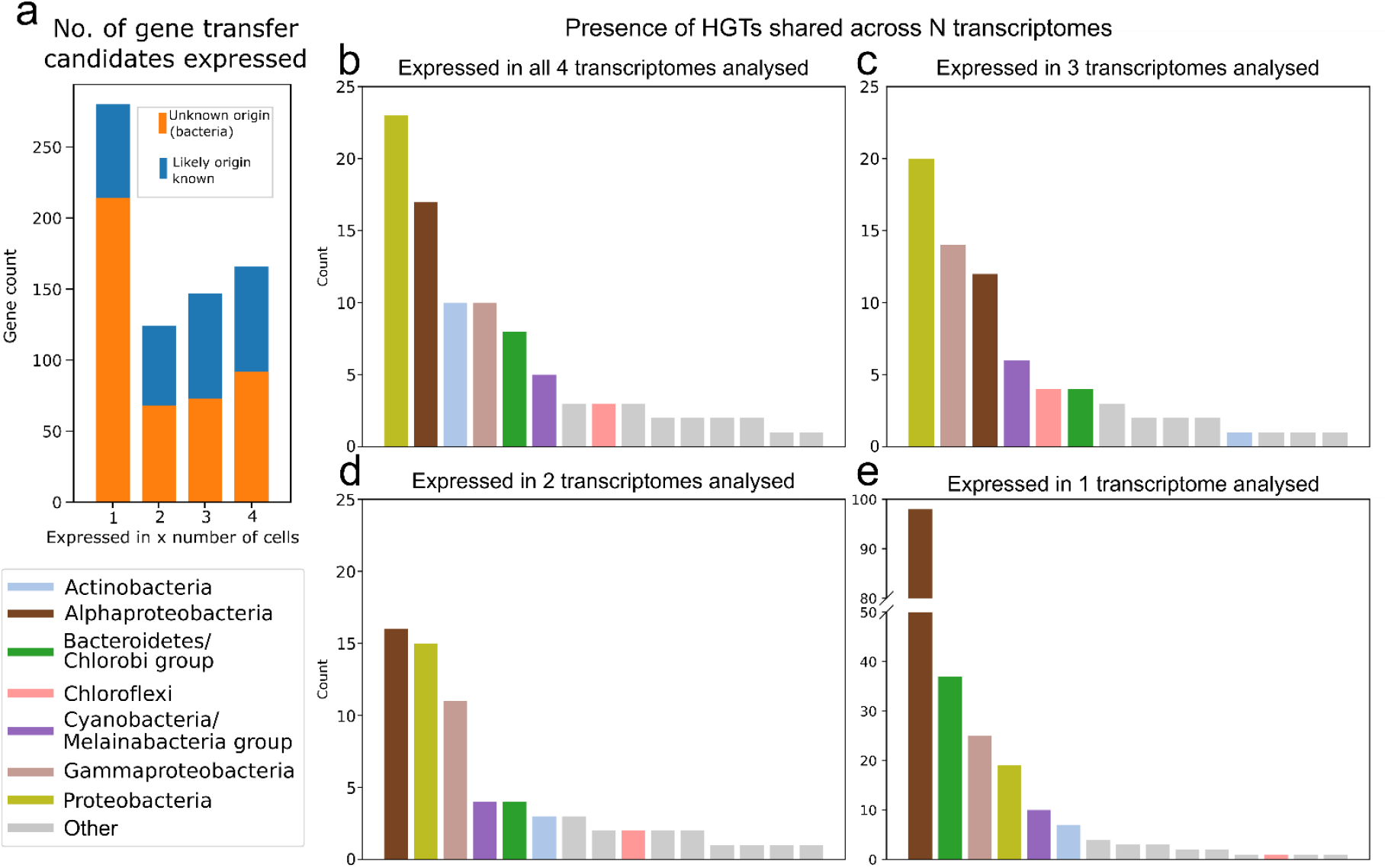
Transcriptome-based analyses of genes of foreign origin in the *Sinophysis* sp. Okigroup1 genome. **a)** Number of candidate bacterial horizontal gene transfers (HGT) identified from the 4 cells sequenced. Candidate HGTs with potential phylogenetic origins are in orange, while those with ambiguous origins are in blue. **b-e)** Presence of HGTs shared across multiple transcriptomes was used to assess reliability and confidence for non-contamination origin. Potential donors of each gene that were expressed in 4, 3, 2, or 1 libraries were used to visualise the diversity of HGTs.

Of the high-confidence HGT candidates, 11 were predicted to be of cyanobacterial origin. Phylogenetic analyses revealed that two were likely to be from endosymbiotic gene transfers, while the others originated from a variety of other cyanobacterial groups. Both EGTs encode for photosynthesis-related genes: C-phycoerythrin beta chain (*pheB* gene) with eukaryotic-like 3’ and 5’ UTRs and expressed across all the sequenced libraries, and a part of the Photosystem II protein D1 (*psbA* gene) with an additional 5’ region consisting of a transmembrane region (Figure S15, Table S4). Interestingly, a HGT candidate encoding the protochlorophyllide reductase B *(porB)* containing a 3’ UTR was also identified. The conserved 5’ dinoflagellate transcript cap, spliced leader, was not identified in most transcripts, including the ones that were predicted to be *bona fide* dinoflagellate transcripts.

### Morphology of *Sinophysis* sp. OKI group 1

Cells are ovoid from the lateral view, with cell sizes ranging from length 47.31-55.62 µm (51.23 ± 1.83 µm, n = 19), width 40.73-48.06 µm (44.23 ± 1.86 µm, n = 19) and a length/width ratio of 1.11-1.24 (1.16 ± 0.033. n = 19) (Figure 1a. S16a, b). Both epitheca and hypotheca are areolate with smooth and round pores with a diameter size 0.166-0.372 μm (0.257 ± 0.042 μm. n = 62)(Figure S16d). Pores are commonly found within areolae and occasionally between areolae. The pores are most prominent on the periphery of the hypotheca, which become sparser closer to the middle of the theca (Figure S16a, c). Epitheca is small and convex, with upturned edges (Figure S16e, f). The plates are asymmetrical, the left plate possesses three anterior projections, one flat projection, and a pair which are protruding (Figure S16e, f). The right plate is slightly overlapped by the hypotheca adjacent to the cingulum (Figure S16f). The cingulum is narrow with list projections from both epitheca and hypotheca (Figure S16c, e, f). The transverse flagellum emerges from the cingulum at the left hypothecate plate. The left hypotheca plate is slightly concave, possessing a deep slit in the centre posterior region (Figure S16a, c). The right hypothetical plate is convex (Figure S16b, c). The sulcus is about 2/3 the length of the right hypotheca plate where the longitudinal flagellum emerges (FigureS16d).

### Description of “Candidatus Geminocystis photosymbiotica” sp. nov

*Geminocystis photosymbiotica* (from photo-for its photosynthetic function, and -symbio, for its symbiotic nature).

Non-motile cyanobacterium from the family Geminocystaceae (order Chroococcales). Spherical in shape with a diameter of 3.95-4.9 pm (4.34 ± 0.23, n = 15), but several individuals are “dumbbell-shaped” with stunted cytokinesis. Cells reside within a double-membraned symbiosome, forming clusters that are arranged at the periphery of *Sinophysis* sp. Okigroupl. The thylakoid membranes are parietally arranged. Cell division by binary fission. Genome size of 2,178,118 bp, and GC content of 34.71%. Complete genome sequence: SAMN48632256. Recognised by oligonucleotide probes: SinoCyanol6S_1, SinoCyanol6S_2 (sequences in materials and methods).

## DISCUSSION

### *Sinophysis* dinoflagellates house two bacterial endosymbionts

Some species from the genus *Sinophysis* have been previously reported to contain ‘green bodies’ hypothesized to be cyanobacterial endosymbionts based on 16S rRNA gene sequencing and fluorescence spectrum emission wavelengths^23^. In *Sinophysis* sp. Oki group 1, we confirm that these represent a single cyanobacterium from the order Chroococcales (sister to G. *herdmanii*), which is phylogenetically distinct from cyanobacterial symbionts from the genus *Synechococcus* harboured by other Dinophysales dinoflagellates^24,25^, indicating their independent acquisition (Figure S5).

Previous observations also showed the presence of a second, much smaller bacterium^23^, which we identified as an alphaproteobacterium from the genus *Nisaea* (Table S1). Volume EM data show that each cyanobacterium is in close contact with multiple alphaproteobacteria and co-localized within the same symbiosome. However, our data do not allow us to confidently determine whether all bacteria are contained in a single symbiosome as reported for other protist-bacteria symbioses^26^. This close association between two symbionts is seen in multiple samples from both our study and the previous report, collected more than 900 km apart^23^. The two symbionts are also predicted to be metabolically coupled, as several pathways, such as the TCA cycle, molybdenum cofactor biosynthesis, and pyrimidine deoxyribonucleotide biosynthesis, which have been lost from the cyanobacterium, are retained by the alphaproteobacterium (Figure 3a, b). Additionally, pantothenate biosynthesis is only complete when combining genes present on the genomes of both the symbionts, resembling other dual symbioses^27-30^.

Given the consistent presence of both symbionts throughout independent sampling sites and time points, and considering the tight association between the two symbionts, the available evidence points to a recent metabolic interdependence (co-obligate) rather than the alphaproteobacterium being a parasite (Table S1). This notion is further supported by the high number of pseudogenes predicted in the alphaproteobacterial genome and short 16S rRNA gene branch length (Figure S3, S6, Table S2). These features are typical of the early stages of an endosymbiotic lifestyle^31^-^32^, suggesting that the alphaproteobacterium was acquired more recently than the cyanobacterium. Such a multi-partner symbiosis is not unique to *inophysis*, as it has also been observed in multiple related dinoflagellates, including Ornithocercus (ectosymbionts) and *Amphisolenia* (endosymbionts). These hosts harbour multiple types of non-photosynthetic bacteria simultaneously with cyanobacteria^21,33^. However, due to their scarce and unculturable nature, whether the potential interactions and their cellular integration level are similar to *Sinophysis* remains unknown.

### The cyanobacterial symbionts are on the way to becoming plastids

Evidence of long-term endosymbiosis of the cyanobacterial symbiont can be observed through the gene losses and pseudogenisation of DNA repair and synthesis, cell membrane/wall structure, and transporters. The loss of assimilatory nitrate reduction and associated pathways indicates a reduction in nitrate uptake requirement or the presence of an alternative method for organic nitrogen acquisition. These losses in the cyanobacterium suggest long-term presence in a stable environment inside the host. However, the cyanobacterium has retained independence in core pathways such as amino acid biosynthesis and glycolysis (Figure 3a).

The cyanobacterium appears to be mostly connected in pairs, forming larger clusters (Figure 4, Movie S1, S2). This is in contrast to the division pattern of free-living *G. herdmanii*, which are separated after division^34^. The presence of multiple semi-divided cyanobacteria in multiple host cells analysed indicates that either they are very actively dividing within the constrained space of the host cell, or the host can block their cell division^35^.

Although established protein transport mechanisms have been repurposed in multiple dinoflagellates with complex plastids^17^, as well as in other symbiotic systems such as *Bigelowiella natans* ^36^ and *P. chomatophora* ^14,37^, our results suggest that they are likely not used for the import of host nuclear-encoded proteins from the *Sinophysis* cytoplasm to the cyanobacteria. Additionally, EGT candidates either lacked identifiable transit peptides or their localisation was predicted to be in other cellular compartments (Table S3). We hypothesize that the import mechanism could be either unique and different from conventional plastid targeting (due to the independent acquisition of their symbiont) or it has not yet been fully established^14,38^.

Cyanobacteria are common nitrogen-fixing symbionts^39,40^. However, in the case of symbiosis with Sinophysis, the cyanobacterium does not contain nitrogen fixation genes and instead functions as a photosymbiont. The production of oxygen measured by nanorespirometry provides strong evidence that the cyanobacterium is photosynthetic (Figure S7). Additionally, the functions of the dinoflagellate EGTs are related to photosynthesis (Figure S15), suggesting that the host can potentially regulate the expression of these genes, promoting photosynthesis on demand. Altogether, the streamlining of the genome, emerging cell division control, and the specific gene transfers to the host point to the cyanobacterium being in an intermediate transition phase to become a primary plastid.

### The role of mixotrophy and co-symbioses in supporting horizontal gene transfer during the early plastid origin

Sinophysis canaliculata has been reported to survive for months in laboratory conditions^41^. However, our attempts to establish its culture in diverse seawater media such as F/2 and Daigo’s IMK, or by feeding it with ciliates *Euplotes vannus* or *E. rariseta* have been unsuccessful. The observed food vacuoles (Figure S10, Movie S1) indicate that they are mixotrophic and would require specific prey for long-term survival. This is not uncommon in dinoflagellates, as many groups are mixotrophic, feeding on eukaryotes and/or bacteria^16,42,43^. The mixotrophic lifestyle likely allows it to be exposed to genetic material from the prey, as proposed for the mixotrophic ancestor of the now fully phototrophic *Paulinella chromatophore*^44−46^, and likely contributes to the diversity of prokaryotic HGTs.

Due to the lack of a high-quality genome of *Sinophysis*, we were unable to analyse eukaryotic HGTs, but the putative bacterial HGT donors (Figure 5a-e) seem to mirror the diversity commonly found in marine benthic ecosystems^47^. The functions of these HGT candidates vary from support to cell lysis. Several HGT candidates are transporters such as ammonium transporters (*amt2*), sulfate transporters (*ybaR*), and other uncharacterized transporters that appear to support the transfer of metabolites, which may contribute to interactions with the symbionts. Other HGT candidates, such as glucose 1-dehydrogenase (*dhgl*) and glutamyl-tRNA aminotransferase subunit A (*gatA*), and choline kinase LicA could be used to promote the growth of the symbiont^48,49^. Additionally, processes related to the photosynthesis of the cyanobacterium could be influenced by genes such as ferrochelatase for haem biosynthesis, and glyoxylate/succinic semialdehyde reductase 1 (*glyrl*), which has been shown to play a role in oxidative stress tolerance^50,51^. On the one hand, protochlorophyllide reductase B (*porB*) has been shown to sustain chlorophyll biosynthesis^52,53^. On the other hand, prophage Rz endopeptidase (*rzpD*) and endolysins may be used for cell lysis^54^.

Only two EGT candidates were identified in the host genome. Both are directly involved in photosynthesis, and their homologs are still present in the cyanobacterial genome. They likely represent new acquisitions, and the released selection pressure on the endosymbiont genes has not yet resulted in their pseudogenization^55-58^. The function of these EGTs supports that during the early stages of the integration, some functions are performed in parallel by both the host and endosymbiont proteins^59^.

### Did heterotrophic dinoflagellates from the genus *Sinophysis* lose their remnant non-photosynthetic plastid after the acquisition of cyanobacterial symbionts?

Although the total loss of the mitochondrial or plastid organelle is relatively rare, it has been previously reported for mitochondria in *Monocercomonides sp*.^60^ and plastids in the parasitic dinoflagellate *Hematodinium sp*.^61^ or the apicomplexan *Cryptosporidium*^62^. A much more common outcome is that, even after losing the organelle’s genome, functionally reduced versions of the organelles are still present and maintained by proteins imported from the host cytoplasm^18^. This scenario has also been hypothesized for many non-photosynthetic dinoflagellates^18,63^. In *Sinophysis*, no non-photosynthetic plastid-like organelles can be identified in our TEM and volume EM data (Figure S9, movie S1). Furthermore, we were unable to detect most genes that are associated with remnant non-photosynthetic plastids in any of our transcriptomes (Figure 5). These results do not rule out the presence of a highly reduced plastid remnant or that the expression of the genes is extremely low or limited to certain life stages. However, since mitochondria-targeted genes were identified (Figure S13), and the transcriptomes show high completeness, it is unlikely that all these genes were missed due to sequencing biases. Assembling the nuclear genome of the uncultivated dinoflagellate and subcellular proteomics data would be required to fully test the absence of the cryptic plastid or its ‘legacy organelle’^64,65^. Importantly, cyanobacterial homologues of these genes are present in the cyanobacterial genome and could be acting as functional replacements (Figure 5), taking over the function of the remnant plastids, making them redundant, and subsequently leading to the loss of the associated genes in the host. This irreversible host-symbiont interdependence has been named the ‘symbiotic rabbit hole’^66,67^.

### What can we learn from the *Sinophysis*-chromatophore system about the early stages of primary plastid endosymbiosis?

Many of the key events that occur during the symbiont-to-organelle transition can already be identified in the *Sinophysis* system. Considering the genome size and gene content of the cyanobacterial symbiont, it is likely that it is in the intermediate stage of transitioning from a symbiont to an organelle^68^. The host seems to already be supporting the symbiont with both native genes and HGTs acquired from various donors, which has been enabled by its mixotrophic lifestyle^6,45,67,69^. On the other hand, the number of EGTs in diverse hosts may be influenced by multiple variables, such as the number and localization of symbionts. In eukaryotes that house a single symbiont per cell, such as in *Angomonas* trypanosomatids and *Braarudosphaera bigelowii* haptophytes^70,71^, only 1 or no EGTs were reported^72,73^, and in *Paulinella* with two symbionts per host, 43 EGT candidates were proposed^74^. In such systems, because of the low number of symbiont cells, lysis of a symbiont cell would be required to access its DNA, which could result either in the loss of the symbiont or in problems with cell division. However, in *Sinophysis*, there are approximately 30-60 symbionts in each cell. The symbiont copy number may thus indirectly influence the EGT frequency. The most highly integrated organelle-like symbionts, such as nitroplasts^71^ or chromatophores, are maintained in only one or two copies per cell^75^, which likely decreases the rate of EGT. Less integrated symbionts, such as cyanobacteria in *Sinophysis*, are maintained at higher loads per cell, potentially due to a less developed host-derived cell division control, which results not only in a lower chance of not passing the symbionts to each daughter cell, but also likely an increased chance for EGT. Future work should thus focus on timing EGT and HGT acquisitions across the *Sinophysis* clade to pinpoint the factors affecting the EGT/HGT ratios during organellogenesis.

In the nitroplast, chromatophore, mitochondria, and plastids of diverse origins, the proteome is a mosaic of both the host and symbiont proteins^14,71,76^. On the other hand, in systems like *Angomonas*-*Kinetoplastibacterium* or diatom-spheroid bodies’, there are few proteins transported to the symbiont^72,77^. In *Sinophysis*, we are experimentally limited by the uncultivated nature of the host. However, we hypothesise that protein targeting to the symbiont is unlikely to be using canonical plastid-import machinery (TOC-TIC translocons). Instead, while host-encoded proteins may still be targeted to the symbiont, their import may rely on novel mechanisms. It is also possible that massive protein import from the host may not be required yet, as the symbiont still retains most essential cellular machineries (Figure 3a). Instead, the symbiont is supported by import of metabolites through a variety of native or HGT transporters (Figure 6) and its close association with the alphaproteobacterial symbiont with complementing metabolic pathways (Figure 3b).

Although symbiont growth is not yet fully controlled, host control of the symbiont seems to start early during the endosymbiont-to-organelle process, likely before protein import establishment. In symbiotic systems where the symbiont is highly integrated, the division of both the host and the symbiont is highly synchronized^78-81^. We noticed that multiple symbiont cells from different *Sinophysis* host cells were found in a mid-division stage before cytokinesis (Figure 4c, 3e, Sll, Movie S2). While not yet fully synchronised, we hypothesise that the host is unable to fully control the growth of the symbiont, but instead uses cell division blocking of the cyanobacterium similar to those found in *Sitophilus-Sodalis*^*82*^ and *Medicago-Rhizobium*^*83*^ systems. As the symbiosis progresses, the host and symbiont divisions will likely become increasingly synchronized similar to other symbionts becoming organelles. Although we did not identify HGTs in the transcriptome that could contribute to division control, host-derived proteins such as dynamin-like proteins could play a role, as seen in *A. deanei*^*35*^.

While most dinoflagellates retain a reduced plastid organelle after losing photosynthesis^18^, the cyanobacterium in *Sinophysis* has likely taken over functions carried out by the remnant plastid (Figure 5). This unexpected result shows that even over short evolutionary timescales, acquiring a new symbiont can lead to the host shifting its metabolic pathways to rely on the symbiont instead of an already established organelle. Such a transition avoids redundancy but effectively locks the interaction into an obligate host symbiont relationship. The ‘shopping bag’ hypothesis is often used to explain the presence of diverse HGTs/EGTs collected via similar gene and symbiont replacements^6,45,67,69^ The intricate evolutionary history of the *Sinophysis* system expands this hypothesis with co-symbioses, symbiont organelle replacements, and massive host gene loss/replacement driven by symbiont turnover (Figure 7). We argue that such a complex symbiotic history should also be considered when discussing the origin of primary plastids.

**Figure 7.**
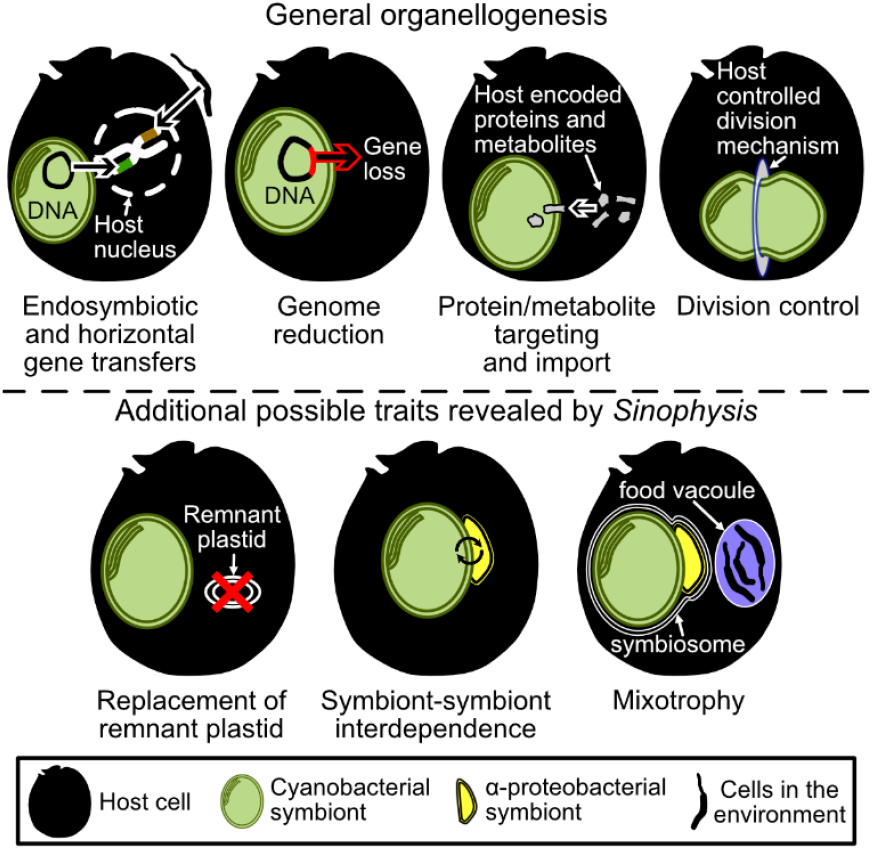
Graphical representation of the key events that lead to plastid organellogenesis. Novel features identified in *Sinophysis* are highlighted below the well-studied features.

## RESOURCE AVAILABILITY

### Lead contact

Further information and requests for resources and reagents should be directed to and will be fulfilled by the lead contact, FH

### Date and code availability

Sequencing data have been deposited under the NCBI BioProject ID PRJNA1256172. All commands used for bioinformatics analyses are available from GitHub: https://github.com/ECBSU/Phua_Sinophysis_2025.git

## ACKNOWLEDGMENTS

This work was supported by the JSPS KAKENHI grant 23K14256 and the HFSP Early Career Grant to FH (RGEC29/2024; https://doi.org/10.52044/HFSP.RGEC292024.pc.gr.194160.) We thank the Scientific Computing (SCDA), Imaging (IMG), and Sequencing (SQC) sections of the Okinawa Institute of Science and Technology for their great support. DL was supported by the JSPS KAKENHI grant 23K05942. This research was partially supported by the Platform Project for Supporting Drug Discovery and Life Science Research from AMED, the Japanese cabinet office and OIST, and a JSPS KAKENHI grant (20K15856) to KCW.

## AUTHOR CONTRIBUTIONS

Conceptualization: PYH, FH, KCW; Field sampling: PYH, FH, KCW, VV; Investigation: PYH, FH, DL, VV, AH, BMH, KCW; Writing - Interpreted data, draft, and approved manuscript: PYH, DL, VV, AH, BMH, KCW, FH; Supervision: FH.

## DECLARATION OF INTERESTS

The authors declare no competing interests.

## DECLARATION OF GENERATIVE AI AND AI-ASSISTED TECHNOLOGIES

The authors used GitHub Copilot and ChatGPT for code auto-completion and debugging. After using the tools, the authors reviewed and edited the content; and take full responsibility for the content of the publication.

## SUPPLEMENTAL INFORMATION

Supplemental information can be found at: 10.6084/m9.figshare.29035775

## METHOD DETAILS

### Key resources table

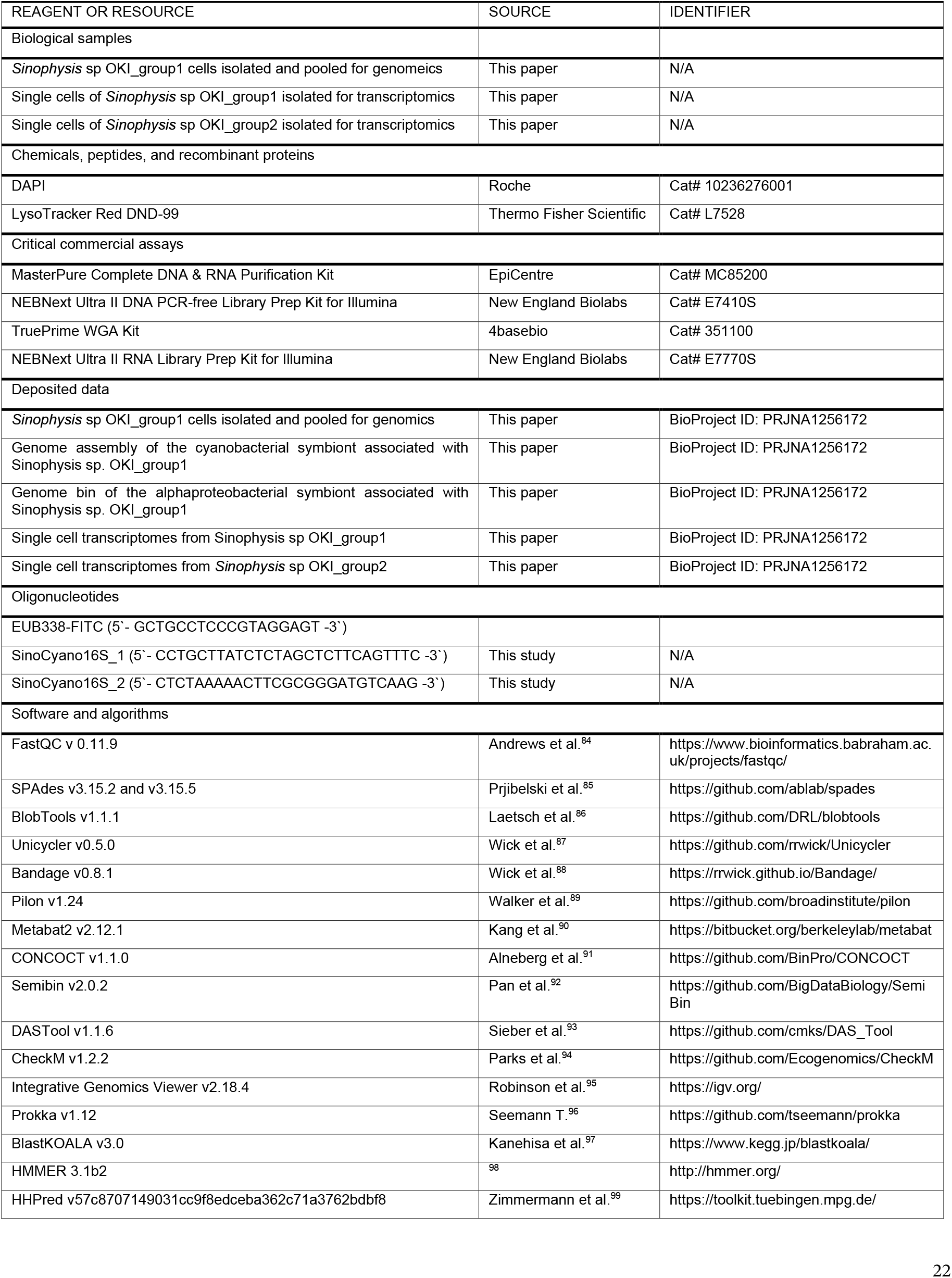
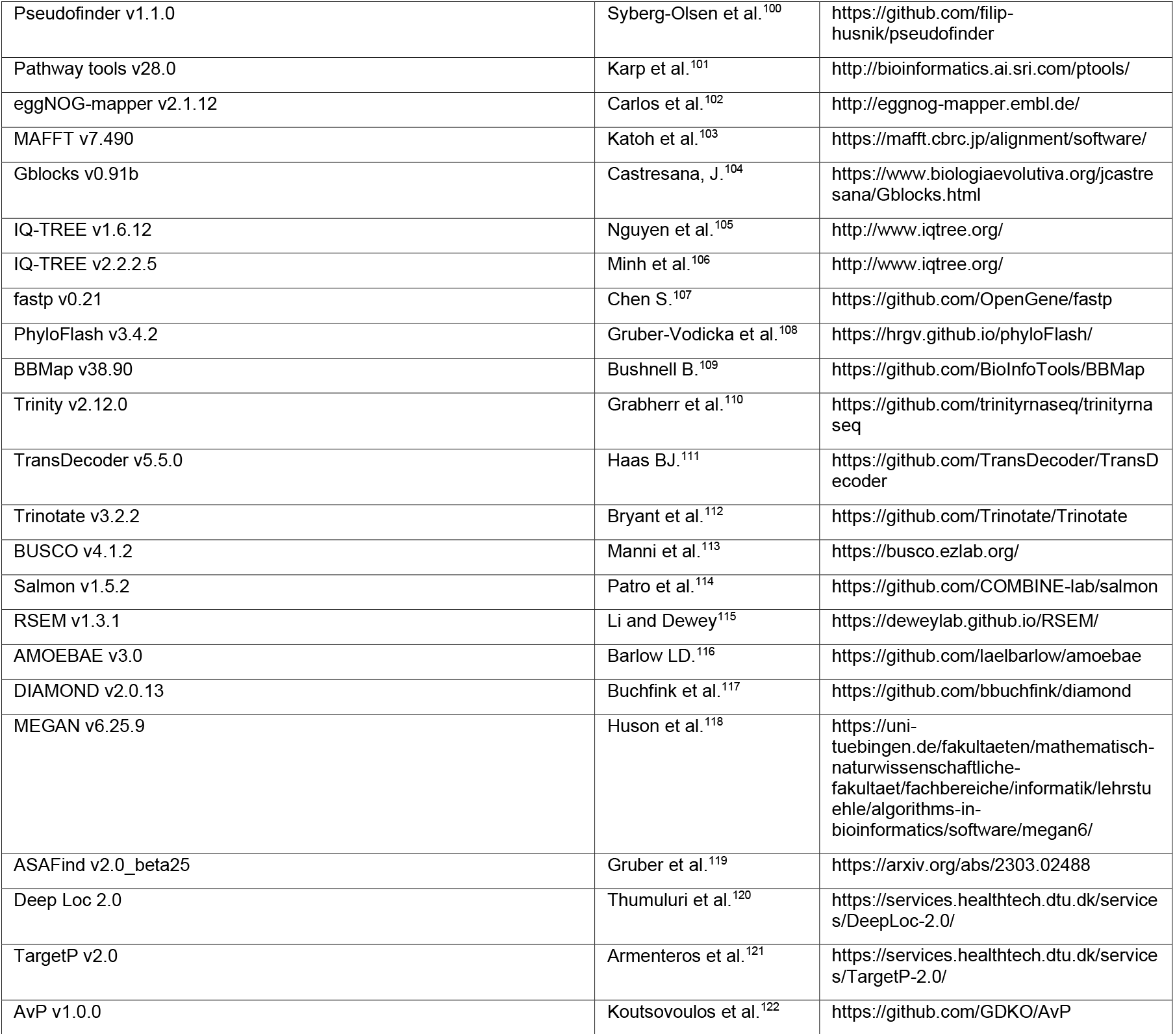

### Sample collection

Crude samples were collected from macroalgae on the coral reefs of Okinawa, Japan (Apogama: 26.4988030°N, 127.8422480°E, and Ikeijima: 26.392003°N, 128.001774°E). Macroalgae were shaken in zip-lock bags filled with seawater and then discarded. Individual cells of *Sinophysis* spp. Okigroupl and Okigroup2 were then isolated from the supernatant under the Olympus CKX41 (Olympus, Tokyo, Japan) and Nikon ECLIPSE Ts2-FL (Nikon, Tokyo, Japan) inverted microscopes with glass pipettes.

### Light and epifluorescence microscopy

For light and epifluorescence microscopy, cells were viewed under the Nikon Ti2-E inverted microscope and imaged using the Nikon Digital Sight 10 camera (Nikon, Tokyo, Japan). For DNA staining, cells were fixed in 4% formaldehyde (FA) for 3 min and stained with DAPI (1 µg/mL) (Roche, Basel, Switzerland). For lysosome staining, cells were stained with a final concentration of 75 nM of LysoTracker Red DND-99 (ThermoFisher Scientific, Massachusetts, United States) and incubated for 2 hr before imaging.

### Fluorescence In Situ Hybridization (FISH) and confocal microscopy

Cyanobacterial symbiont-specific 16S rRNA-specific probes were designed using Mesquite v3.81 ^123^ after aligning the sequences obtained from this study with the SILVA database. The specificity of the probe sequences was then checked with TestProbe 3.0^124^. Two regions were selected, and probes were labelled at the 5’ end with the Alexa 405 fluorophore (ThermoFisher Scientific, Massachusetts, United States) (Table S5). Fresh samples were collected using glass pipettes and fixed in 4% FA at room temperature for 1 hr. The cells were then transferred into wells with 70% EtOH and kept at −20°C. Permeabilization was performed using 100% acetone for 30 min, then washed with PBST (0.3% v/v Triton X-100) 3 times. The cells were then washed with hybridisation buffer (0.9M NaCl, 0.01% SDS, 20mM Tris HCl pH 7.2) 3 times. The hybridisation was performed with 0.1 µM probe at 46 °C for 2 hr. Cells were then washed with PBST 3 times and mounted with ProLong Diamond Antifade Mountant (ThermoFisher Scientific, Massachusetts, United States) on glass slides and imaged using a ZEISS Axio Observer with LSM 980 confocal microscope.

### Transmission Electron Microscopy (TEM)

Cells were fixed in 2.5% glutaraldehyde (FUJIFILM Waka, Osaka, Japan) for 30 min, washed with seawater 3 times for 5 min each, and post-fixed with 1.5% osmium tetroxide (Nisshin-EM, Tokyo, Japan) for 1 hr. The cells were washed with 0.2 µm filtered seawater 3 times and then with **MiliQ** 3 times for 5 min. They were then dehydrated with a stepwise series of EtOH (60%, 70%, 80%, 90%, 100%, 100%, 100%) for 5 min each step. The cells were then permeabilised with a 50% acetone and EtOH mixture for 5 min, 100% acetone twice for 5 min, and then incubated in a 2:1 acetone and low viscosity epoxy resin (Agar Scientific, Rotherham, United Kingdom) mixture for 2 hr at room temperature. The mixture was exchanged with 100% resin twice and then left overnight at RT. The polymerisation was done at 70 °C for 30 hr and ultrathin sections 50 nm thick were cut using an ultra 45° diamond knife (DiATOME, Nidau, Switzerland) and mounted on formvar (SPI supplies, West Chester, USA) coated slot grids. Sections were imaged with Hitachi-7400 (Hitachi, Tokyo, Japan) and JEM-1400 Flash (JEOL, Tokyo, Japan) transmission electron microscopes, equipped with AMT C4742-57-12ER (Advanced Microscopy Techniques, Massachusetts, USA) and Hamamatsu ORCA-Flash4.0 (Hamamatsu, Shizuoka, Japan) cameras, respectively.

### Focused Ion Beam Scanning Electron Microscopy (FIB-SEM)

Cells were fixed in 2.5% glutaraldehyde for 30 min, washed with 0.2 µm filtered seawater 3 times for 5 min each, and post-fixed with 2% osmium tetroxide and 1.5% potassium ferrocyanide (ScyTek Laboratories, Utah, USA) for 1 hr. They were then stained *en-bloc* in 2% uranyl acetate overnight. To remove uranyl acetate, they were washed with filtered seawater 3 times and then with MiliQ 3 times for 5 min each wash. Dehydration was carried out with a stepwise increase of EtOH concentration (70%, 80%, 90%, 3x 100%) for 5 min each step. The cells were permeabilised with 50% acetone and EtOH mixture for 5 min, with 100% acetone twice for 5 min, and in 2:1 acetone and low viscosity epoxy resin mixture for 2 hr. The mixture was exchanged with 100% resin twice and then left overnight at room temperature. Polymerization was done at 70 °C for 30 hr. Then 50 nm ultrathin sections on formvar-coated slot grids were used to check fixation quality on a Jeol JEM-1230R Transmission Electron Microscope (JEOL, Tokyo, Japan). The samples were imaged as follows: 20 nm slices were milled with Ga ions at an acceleration voltage of 30 kV and a current of 2.4 nA using a Helios NanoLab 650 FIB system (FEI, Eindhoven, The Netherlands). The remaining block-face was imaged with an electron current of 800 pA at 1.5 keV acceleration voltage at a dwell time of 6 µs using the through-the-lens backscatter-electron detector (TLD-BSE). The frame size was 6144 x 4096 pixels with a horizontal field of view of 61.44 µm, resulting in a pixel size of 10 nm.

### FIB-SEM data segmentation

Cyanobacteria, food vacuoles, nucleus and theca were segmented in Microscopy Image Browser v2.9010^125^. Cyanobacteria were segmented using the graphcut algorithm on a watershed-based processing of supervoxels. The number of supervoxels and the coefficient of edge values were adjusted based on location and sensitivity required. The food vacuoles, and nucleus were segmented using manual segmentation and interpolated. The theca was segmented using black/white thresholding combined with manual segmentation. Mitochondria and alphaproteobacteria were segmented with ilastik vl.4.0^126^ using a two-stage autocontext workflow. For both stages of training, all features within the 0-4 sigma range were computed in 3D and used. Probabilities from stage two prediction were exported to Microscopy Image Browser for manual clean-up. Volumes of the generated models were calculated in 3D using MIB. Contours were exported to IMOD 3D v5.1^127^ for meshing and rendering was performed using Blender v3.5.

### Genome sequencing

Cells were isolated from fresh samples, washed twice in filtered seawater, and then transferred to a 1.5 mL Eppendorf tube with seawater using glass pipettes. DNA was extracted using the MasterPure Complete DNA & RNA Purification Kit (EpiCentre, Madison, USA) following the manufacturer’s protocol from approximately 400 pooled cells. The purity and concentration of DNA were checked on a Qubit fluorometer and a NanoDrop 1000 spectrophotometer (Thermo Fisher Scientific, Massachusetts, United States), respectively. PCR-free shotgun sequencing libraries were prepared using the NEBNext Ultra II kit (New England Biolabs, Massachusetts, USA) following the manufacturer’s protocol. Multiplexed libraries were pooled and sequenced on an Illumina NovaSeq 6000 platform with 2 x 150 bp paired-end reads.

For long-read sequencing, the DNA was amplified using the TruePrime WGA Kit (4basebio, Cambridge, United Kingdom) following the manufacturer’s protocol and then debranched using the T7 endonuclease (New England Biolabs, Massachusetts, USA). Sample quality was measured using FEMTO pulse (Agilent, California, USA), and the short read fragment buffer was used during the 1D Ligation library preparation. Libraries were loaded on an Rl0.4 flow cell for long-read sequencing using Oxford Nanopore MinION (Oxford Nanopore Technologies, Oxford, United Kingdom). All sequencing runs were performed at the Sequencing Section ofOIST.

### Genome assemblies and analyses

Sequence quality was assessed with FastQC v 0.11.9^84^. Total metagenome assembly from short reads was performed using SPAdes v3.15.2^85^ with the metagenome settings. Symbiont contigs were selected and extracted using BlobTools vl.1.1^86^. Additionally, contigs that had been identified to be cyanobacterial (>90% identity) were also identified using the NCBI BlastN search against the nt database. Reads that mapped to the symbiont and cyanobacterial contigs were extracted using bamfilter module from BlobTools. Hybrid reassembly with only the cyanobacteria short reads and all long reads was performed using unicycler v0.5.0^87^ with a minimum bridge quality set to 5. The assembly graph was examined in Bandage v0.8.1^88^ and closed manually. From the assembly graph, a region with higher coverage (4-lOx) than the rest of the genome was inferred as a plasmid. The closed genome and plasmids were then polished with Pilon vl.24^89^. The genome underwent 23 Pilon iterations, while plasmids underwent 7 and 11 iterations. The alphaproteobacterial symbiont bin was extracted from the hybrid reassembly generated by SPAdes v3.15.5 with the metagenome setting. The binning pipeline consisted of Metabat2 v2.12.1^90^, CONCOCT vl.1.0^9^1,Semibin v2.0.2^92^, then refined with DASTool vl.1.6^93^. The assessment of genome completeness and contamination was performed using CheckM vl.2.2^94^. The contig containing the 16S rRNA gene, partial sequence, was identified from the assembly graph using Bandage v0.8.1 and manually included in the bin. Reads mapped on the genome were then inspected using IGV v2.18.4^95^ before and after polishing with Pilon vl.24 with for 24 iterations.

Annotation was performed using Prokka vl.12^96^, and BlastKOALA v3.0^97^. The possible function of the hypothetical proteins from the cyanobacteria plasmid was further annotated with HMMER v3.lb2^98^ against the Pfam database (RELEASE 37.4) and HHpred9^9^ online server using default parameters against the PDB_mmCIF70_25_may database. Pseudogenes were identified using Pseudofinder vl.1.0^100^ against *G. herdmanii* and *N. acidiphila* as references for the cyanobacterial and alphaproteobacterial genome, respectively. The alphaproteobacterial genomes before polishing had fewer pseudogenes and thus was used for downstream analysis. Metabolic pathways were predicted and visualised in the KEGG reconstruct tool^128^ and Pathway tools v28.0^101^. COG annotations were predicted by the eggNOG-mapper v2.1.12 web server^102^ against the eggNOG 5 database. A circular map of the genome with the gene and CG content was generated using the Proksee web server^129^. For proteins that were present in G. *herdmanii* but absent in *G. photosymbiotica*, the absence was confirmed by using Blastx against the total assembly. From these, cyanobacterial contigs were extracted and identified using BlastX web server against the nr database, restricting the search to cyanobacteria.

### Phylogenetics

The 16S rRNA gene sequences from the two symbionts and the 18S rRNA gene from *Sinophysis* were aligned within their respective groups using the MAFFT v7.490 algorithm^103^ together with sequences retrieved from the NCBI nt database. Alignments for the cyanobacteria were performed with the automatic setting, while *Nisaea* and *Sinophysis were* aligned with the L-INS-i algorithm. For the *Nisaea* alignment, the overhanging ends of the longest sequences were manually trimmed to match the rest of the alignment. The cyanobacterial and *Sinophysis* alignments were trimmed using Gblocks v0.9lb^104^ with default block parameters but allowing for positions with up to 50% gaps (i.e., half allowed gap position setting). For cyanobacteria, IQ-TREE vl.6.12^105^ was used to select the TVMe+I+G4 substitution model according to the Bayesian Information Criterion and to infer the Maximum Likelihood (ML) phylogenetic trees with 1,000 bootstrap pseudo-replicates. For *Nisaea* and *Sinophysis*, the substitution models TN+F+I+G4 and TIM2+ F+R2 were selected, and ML phylogenetic trees were inferred using IQ-TREE V2.2.2.5^106^ with 1,000 bootstrap pseudoreplicates.

### Transcriptome sequencing

Individual cells were manually lysed using a glass pipette in 0.2 mL thin-walled PCR tubes. Single-cell cDNA samples were prepared following the Smart-seq2 protocol, except that the Superscript II Reverse Transcriptase was used, and a ligation-based library preparation kit was used instead of Nextera XT^130^. Eighteen PCR-plus sequencing libraries were prepared using the NEBNext Ultra II kit (New England Biolabs, Massachusetts, USA) following the manufacturer’s protocol (with 6 amplification cycles). The libraries were multiplexed, pooled, and ten libraries were sequenced on the Illumina NovaSeq 6000 platform, the remaining eight on the Illumina NovaSeqX platform, with 2 x 150 bp paired-end reads.

### Transcriptome assembly and analyses

Sequence quality was assessed with FastQC v 0.11.9^84^. Sequencing and Smart-seq2 adaptors were removed using fastp v0.21^107^ and their removal again confirmed using fastqc v0.11.9^84^. Then PhyloFlash v3.4.2^108^ was used to check for potential contamination and retrieve rRNA gene sequences of all organisms present in the data. Raw reads of the cyanobacterial symbiont were filtered out using BBSplit v38.90^109^ with the default minimum mapping identity setting. The reads from Oki group 1 identified by 18S rRNA genes were extracted and aligned against other *Sinophysis* spp. 18S rRNA genes retrieved from NCBI nt database using MAFFT v7.490. The alignment was trimmed with Gblocks with default settings but allowing for up to 50% gap positions, and a maximum likelihood tree was generated using the TIM2+F+R2 substitution model on IQ-TREE V2.2.2.5. The reads were pooled and assembled *de nova* using Trinity v2.12.0^110^, and protein coding genes were predicted using TransDecoder vS.5.0^111^ and annotated in Trinotate v3.2.2^112^. Transcripts from the alphaproteobacterial symbiont filtered out by identification using NCBI BlastN searches (>98% identity, lO0bp length) against the symbiont bin, and removed from the assembly. Completeness of the dinoflagellate transcriptomes was checked using BUSCO v4.l.2^113^ against the alveolata_oblO lineage dataset. Transcript quantification was performed using salmon vl.5.2^114^ and RSEM^115^ with the bowtie alignment method. The presence of genes typically found in non photosynthetic plastids was identified using the AMOEBAE v3.0^116^. Transcripts that were classified as of bacterial origin (i.e., likely contaminants) were identified using Diamond v2.0.13^117^ BlastP against the nr database and MEGAN v6.25.9^131^, and removed from the AMOEBAE results. The presence of signal peptides and protein localization was predicted using ASAFind v2.0_beta25^119^, Deep Loe 2.0^120^, and TargetP v2.0^121^. The dinoflagellate spliced leaders^132^ were queried against the transcriptome using NCBI BlastN with the ‘short queries’ option.

To identify horizontal gene transfer candidates, the taxonomic placement of each protein was identified using Diamond v2.0.13 BlastP with the sensitive option against the NCBI nr database. The taxonomy classes of each sequence were identified using MEGAN v6.25.9 with meganMapDB-Feb2022 database, and bacterial sequences were classified as HGTs. Genes that were expressed in at least 3 of the samples analysed were considered to be less likely to be from contamination. Additionally, the presence of longer eukaryotic-like UTRs was used as an indicator of eukaryotic origin. Proteins that were identified to be of prokaryotic origin were then screened again using AvP vl.0.0^122^. From the collection of HGTs, EGT candidates were identified by BlastP against the protein-coding genes predicted from the cyanobacterial symbiont. The sequences of EGT candidates were aligned against proteins obtained from the NCBI nr database using MAFFT v7.490 aligner employing the L-INS-i algorithm. The results were, trimmed with Gblocks v0.91b using default block parameters, but allowing for positions with up to 50% gaps. Substitution models were selected using IQ-TREE v2.2.2.5 for the C-phycoerythrin beta chain and photosystem II protein Dl genes, which were Q.insect+R3 and LG+R3, respectively. and phylogenetic trees were inferred with 1,000 bootstrap pseudo-replicates. Protein functions of the EGT candidates were predicted with InterPro webserver^133^.

The metabolic pathway completeness for the host transcriptome, symbiont genomes, as well as free-living reference genomes were predicted using a custom pipeline. The pipeline performs functional annotation using eggNOG-mapper v2102 and converts the generated KEGG annotation terms into KEGG pathway completeness values (https://github.com/ECBSU/genomics-scripts/tree/main/Functionalannotation/KEGGstand_in_house_KEGG_annotation).

### Oxygen microsensor measurements

Freshly sampled cells were placed at the bottom of a microtube (500 µm inner diameter) filled with microfiltered natural seawater. Oxygen gradients in the 1 mm above the cells were measured with 100 µm step size for each measurement, using a 50 µm tip diameter Clark-type microelectrode (Unisense, Denmark) fixed on a motorized micromanipulator (Unisense, Denmark). Before the measurements, the microelectrode was 2-point calibrated using a 10 g/L sodium hydrosulfite solution (Sigma-Aldrich, Missouri, United States) and aerated seawater. Measurements were performed under decreasing light intensity (at 300, 115 and O µmol*O*_2_/m2/s photosynthetically active radiations provided by a video-grade LED panel and controlled by a MQ-200 quantum meter, Apogee instruments). Individual photosynthesis rates were calculated using Fick’s first law of diffusion following^134,135^. The calculation pipeline and raw data are publicly accessible at https://github.com/ECBSU/nanorespirometry.

A total of four replicates were measured for *Sinophysis* (3 replicates containing 3 cells and 1 replicate containing 1 cell), one replicate was measured for *Prorocentrum* (using 16 cells) and was considered as a positive control, and one blank (containing only filtered seawater) was used, showing low oxygen production (< 22 pmol*O*_2_/microtube/day). All the measurements were performed in a temperature-controlled room (with measured water temperature ranging from 23.0°C to 23.4°C).

## Supplemental information

**Figure S1.**
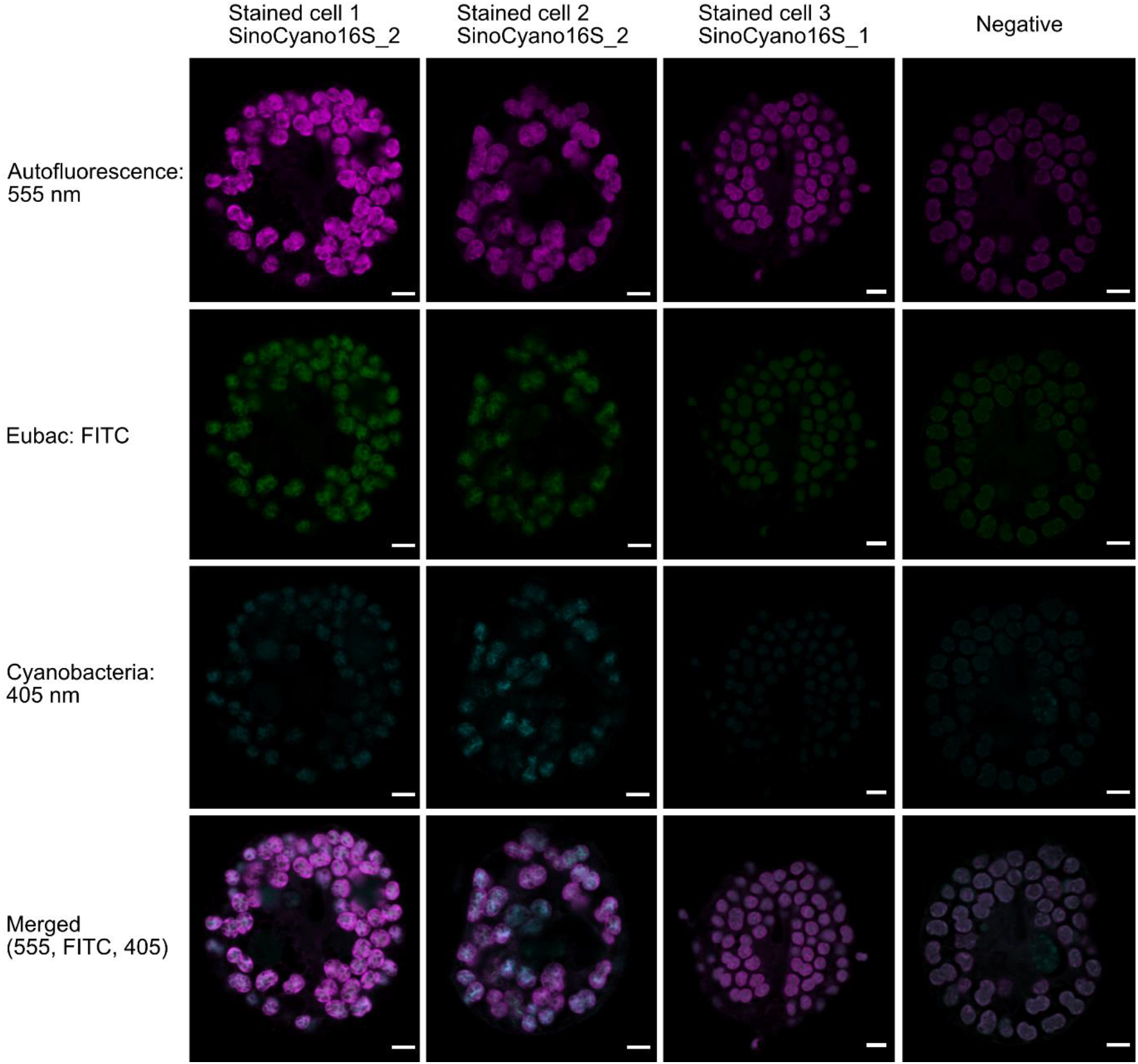
Confocal micrographs of FISH-stained *Sinophysis* sp. cells. Cells stained with 16S rRNA gene probes, symbiont-specific probes (SinoCyano16S_2 and SinoCyano16S_1), and general eubacterial probes (Eubac-FITC). The negative cell was used as an autoluorescence and no-probe control. Autofluorescence (magenta), EUB338-FITC (green), SinoCyano16S_1 and SinoCyano16S_2 probes (cyan). Scale bar = 5 µm

**Figure S2.**
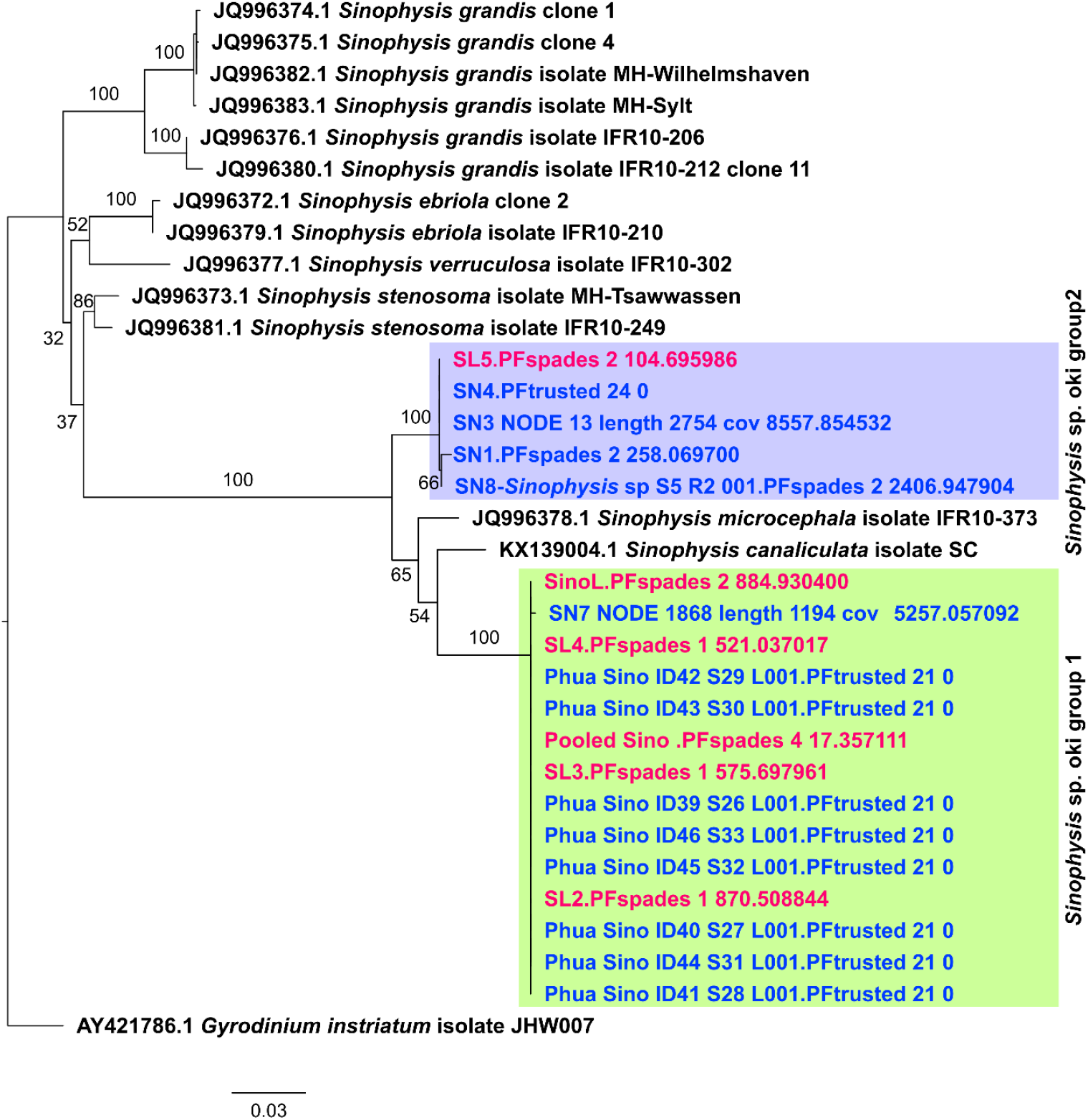
Maximum likelihood phylogenetic tree inferred from the 18S rRNA gene sequences of *Sinophysis* spp newly acquired in this study. The tree was inferred under the TIM2+F+R2 substitution model using 1,000 bootstrap pseudo-replicates. Samples were collected from Apogama (red) and Ikeijima (blue) in Okinawa. The sequences form two clusters, likely two new species, which we refer to as group 1 and group 2 throughout the manuscript.

**Figure S3.**
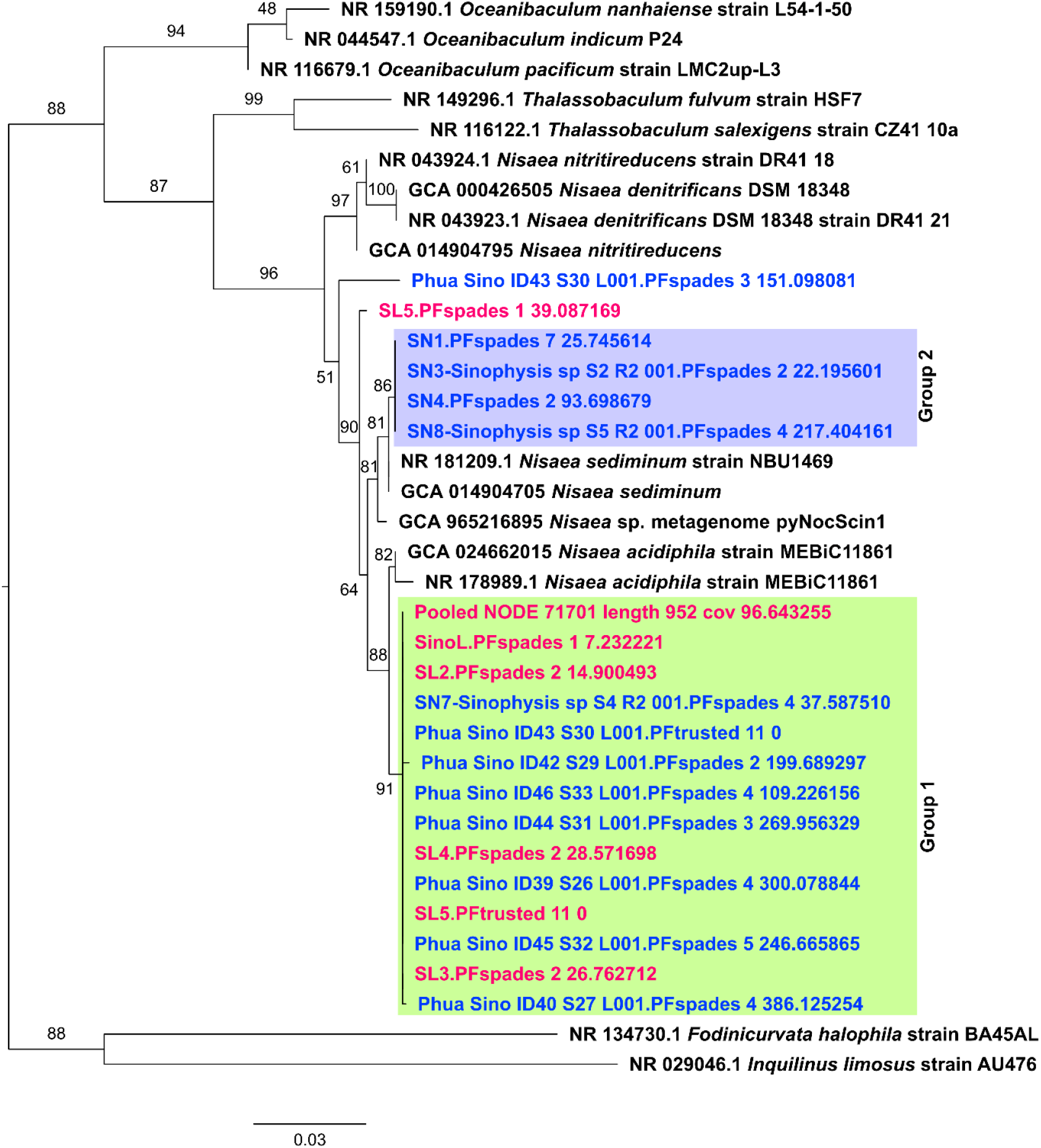
Maximum likelihood phylogenetic tree inferred from the 16S rRNA gene sequences from *Nisaea* spp. The tree was inferred under the TN+F+I+G4 substitution model using 1,000 bootstrap pseudo-replicates. Samples were collected from Apogama (red) and Ikeijima (blue) in Okinawa.

**Figure S4.**
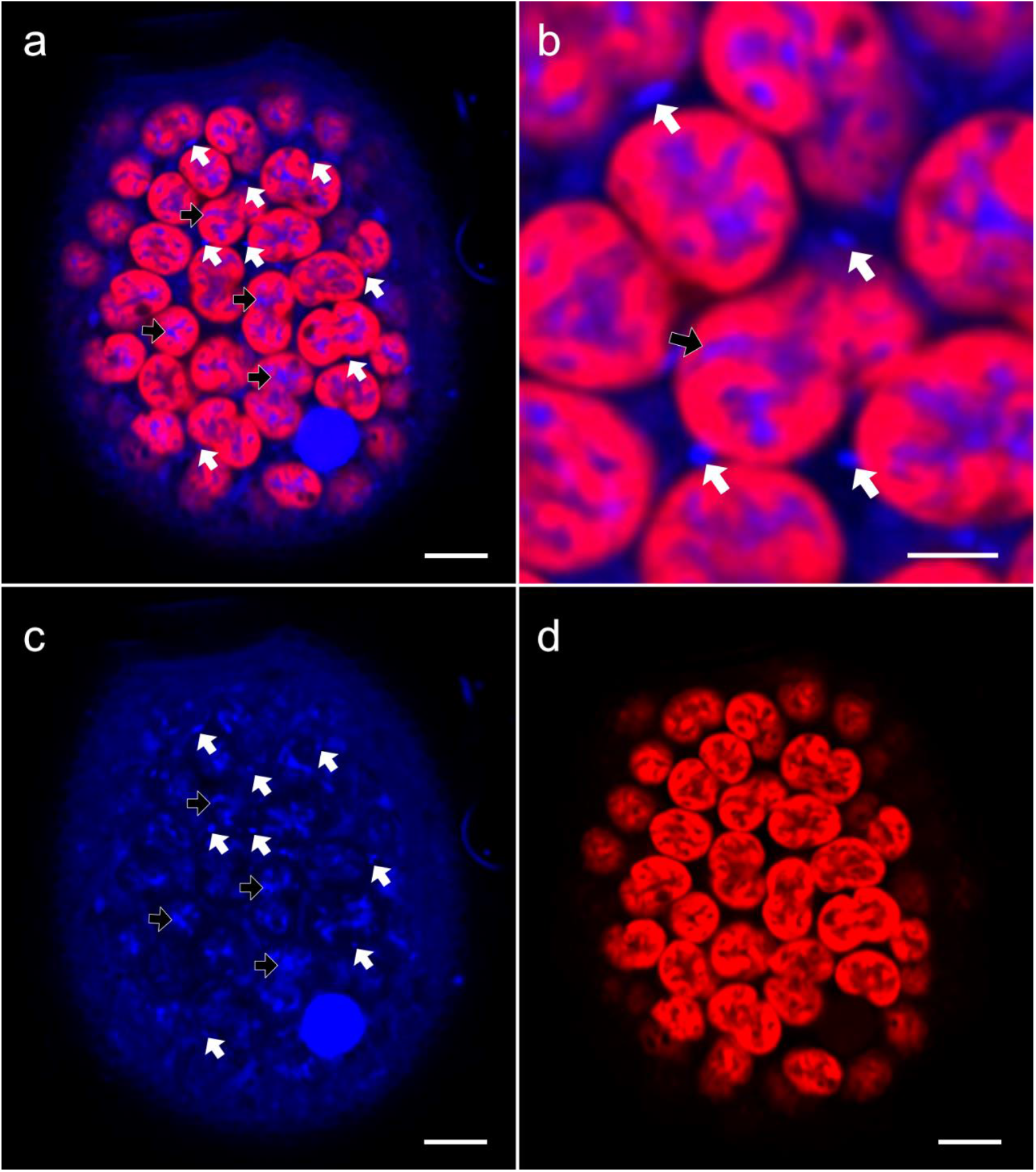
Confocal micrograph of DAPI-stained *Sinophysis* sp. cells. **a, b)** Merged DAPI and Cy5 (autofluorescence) channels with signal localised within the cyanobacterial symbionts (black arrows) and *Nisaea* (white arrows). **c)** DAPI channel (blue, excitation λ: 405 nm), **d)** Cy5 channel (red, excitation λ: 633 nm).

**Figure S5.**
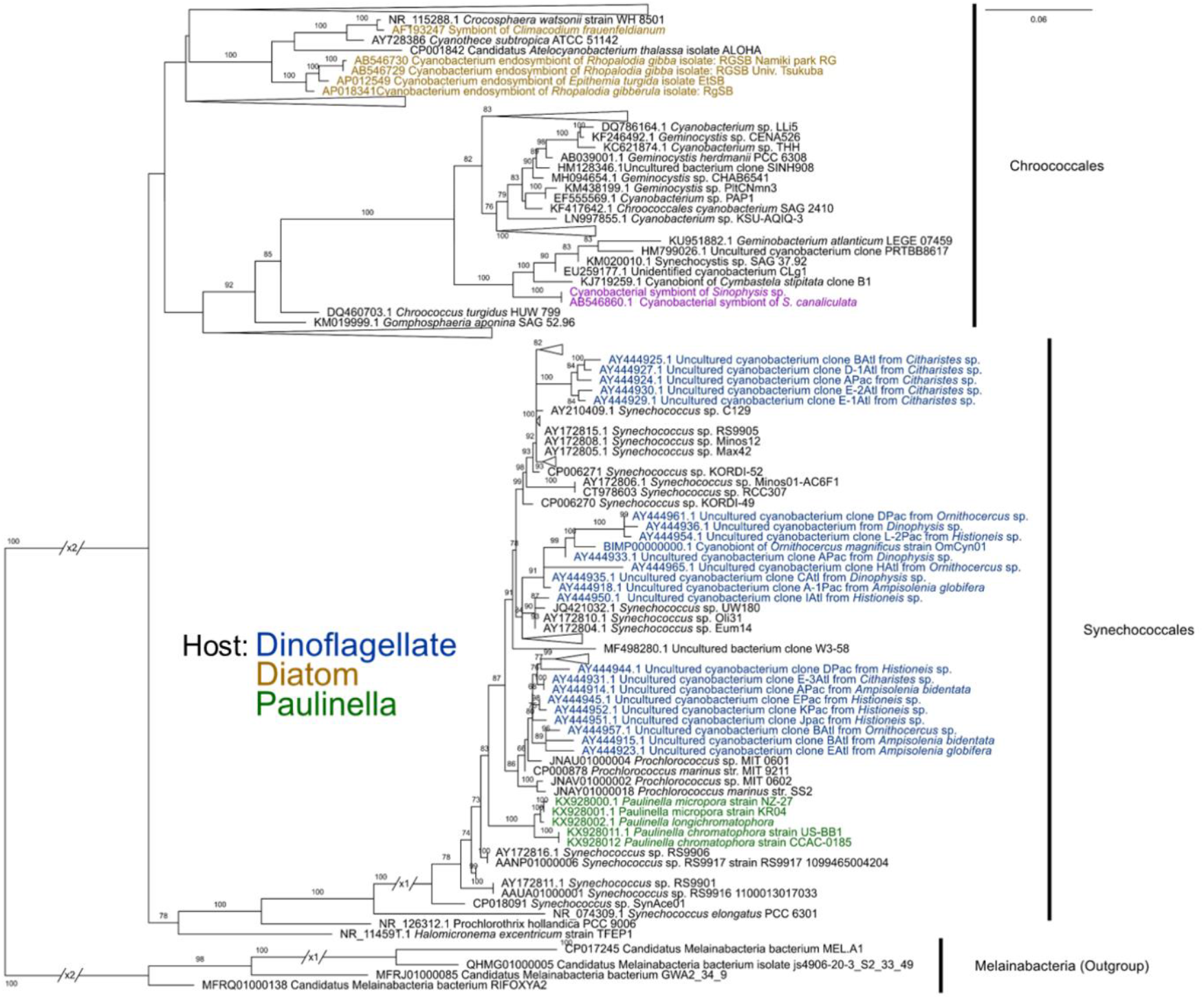
Maximum likelihood phylogenetic tree of 16S rRNA gene sequences from representative Cyanobacteria. The tree was inferred under the TVMe+I+G4 substitution model using 1,000 ultrafast bootstrap pseudo-replicates. Cyanobacteria that are known to be symbionts of microalgae are coloured by the host: dinoflagellate (blue), diatom (brown), *Paulinella* (green), and *Sinophysis* (purple).

**Figure S6.**
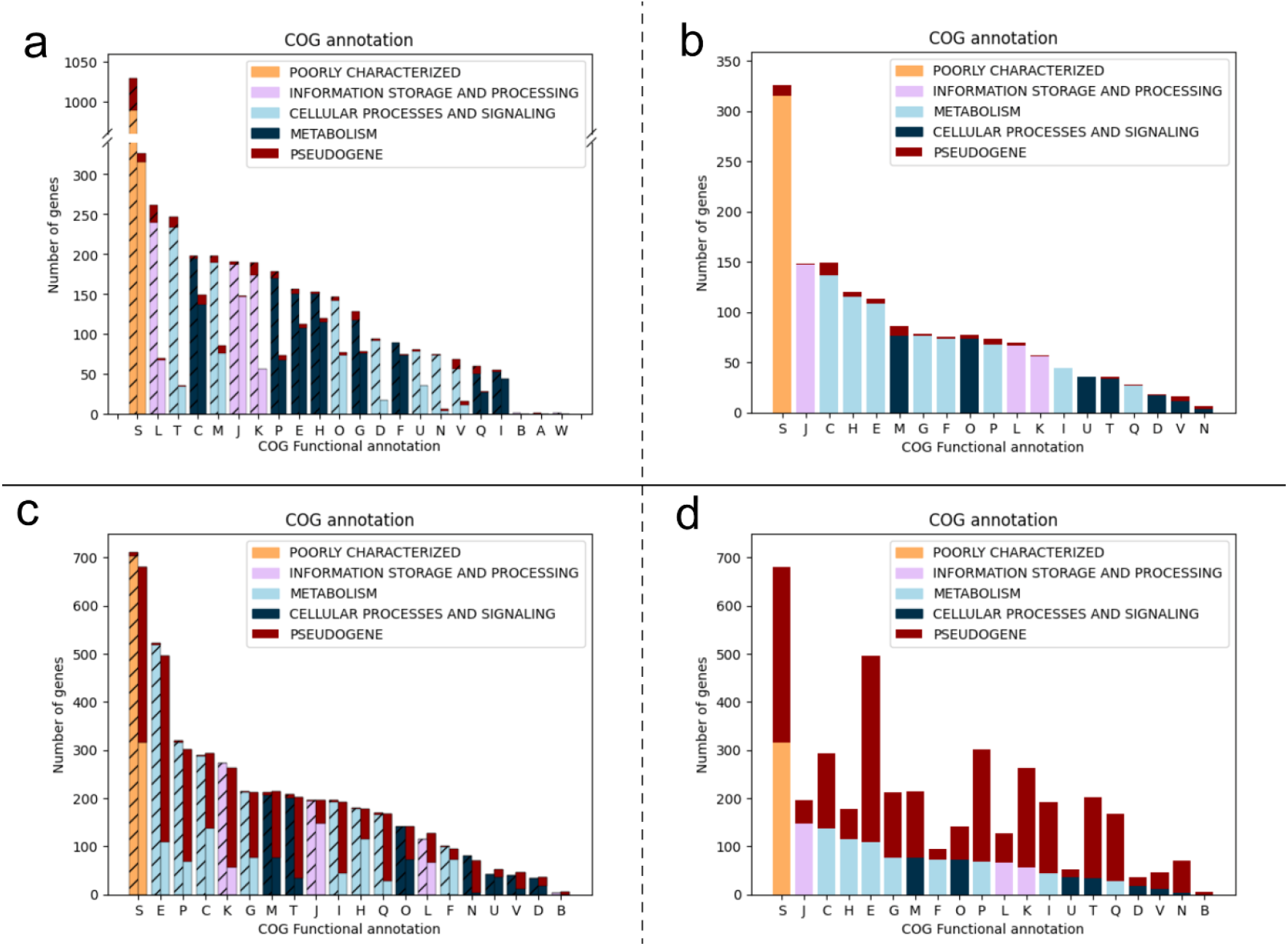
Comparison of COG annotations for genes present in the symbiont genomes. **a, b)** Genes present in the cyanobacterial (top), and **c, d)** alphaproteobacterial symbionts (bottom), as compared against their closest free-living relatives. (hatched bars). COG categories: [J] Translation, ribosomal structure and biogenesis, [A] RNA processing and modification, [K] Transcription, [L] Replication, recombination and repair, [B] Chromatin structure and dynamics, [D] Cell cycle control, cell division, chromosome partitioning, [Y] Nuclear structure, [V] Defense mechanisms, [T] Signal transduction mechanisms, [M] Cell wall/membrane/envelope biogenesis, [N] Cell motility, [Z] Cytoskeleton, [W] Extracellular structures, [U] Intracellular trafficking, secretion, and vesicular transport, [O] Posttranslational modification, protein turnover, chaperones, [X] nan, [C] Energy production and conversion, [G] Carbohydrate transport and metabolism, [E] Amino acid transport and metabolism, [F] Nucleotide transport and metabolism, [H] Coenzyme transport and metabolism, [I] Lipid transport and metabolism, [P] Inorganic ion transport and metabolism, [Q] Secondary metabolites biosynthesis, transport and catabolism, [R] General function prediction only, [S] Function unknown

**Figure S7.**
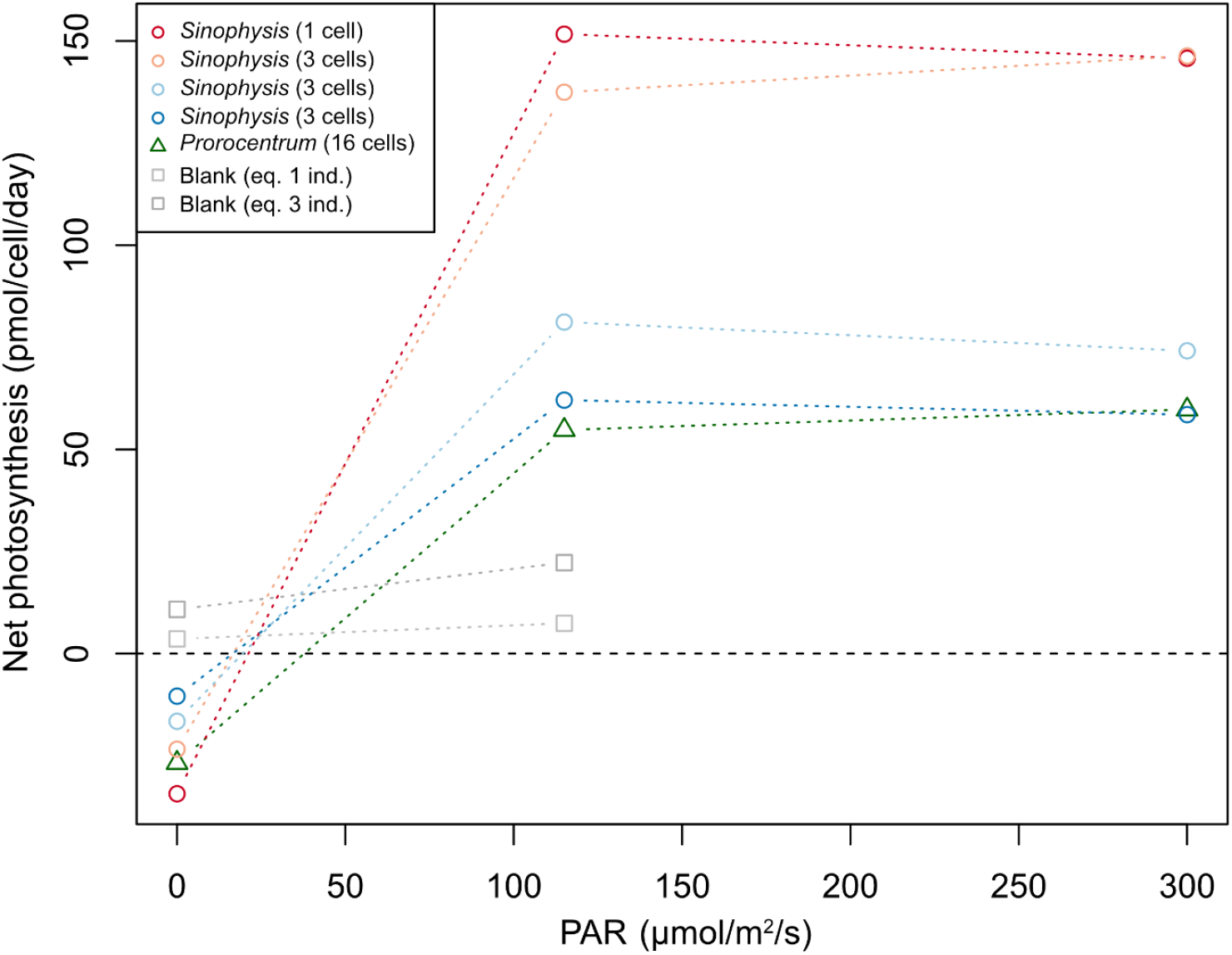
Relationship between net photosynthesis and photosynthetically active radiation (PAR) in *Sinophysis* sp.

**Figure S8.**
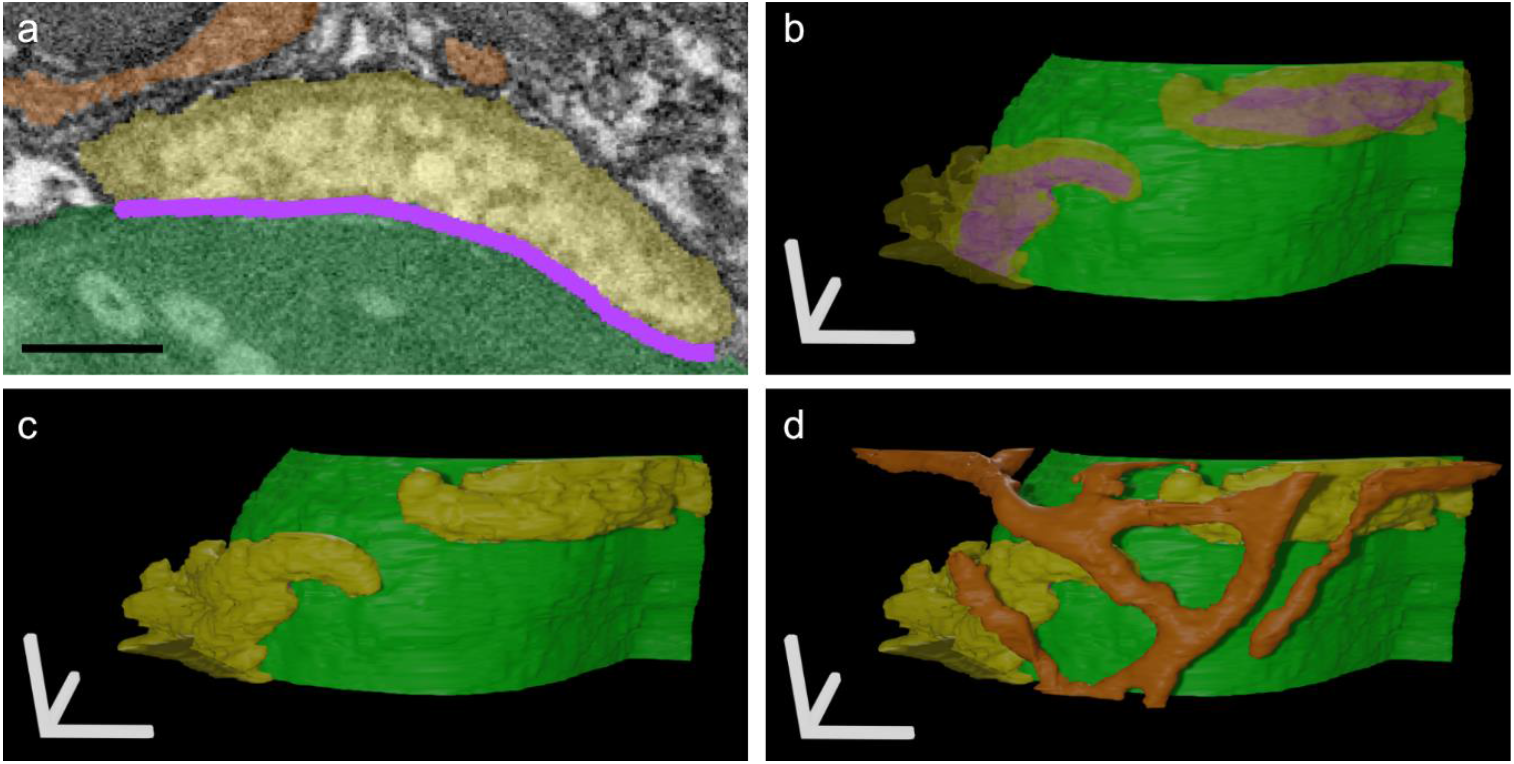
Reconstruction of a FIB-SEM micrograph of symbionts in contact with each other. **a, b, c)** Contact sites (purple) between a cyanobacterium (green) and alphaproteobacteria (yellow). **d)** Mitochondria (orange) are not in contact. Scale: **a)** 500 nm, **b - d)** 1 μm.

**Figure S9.**
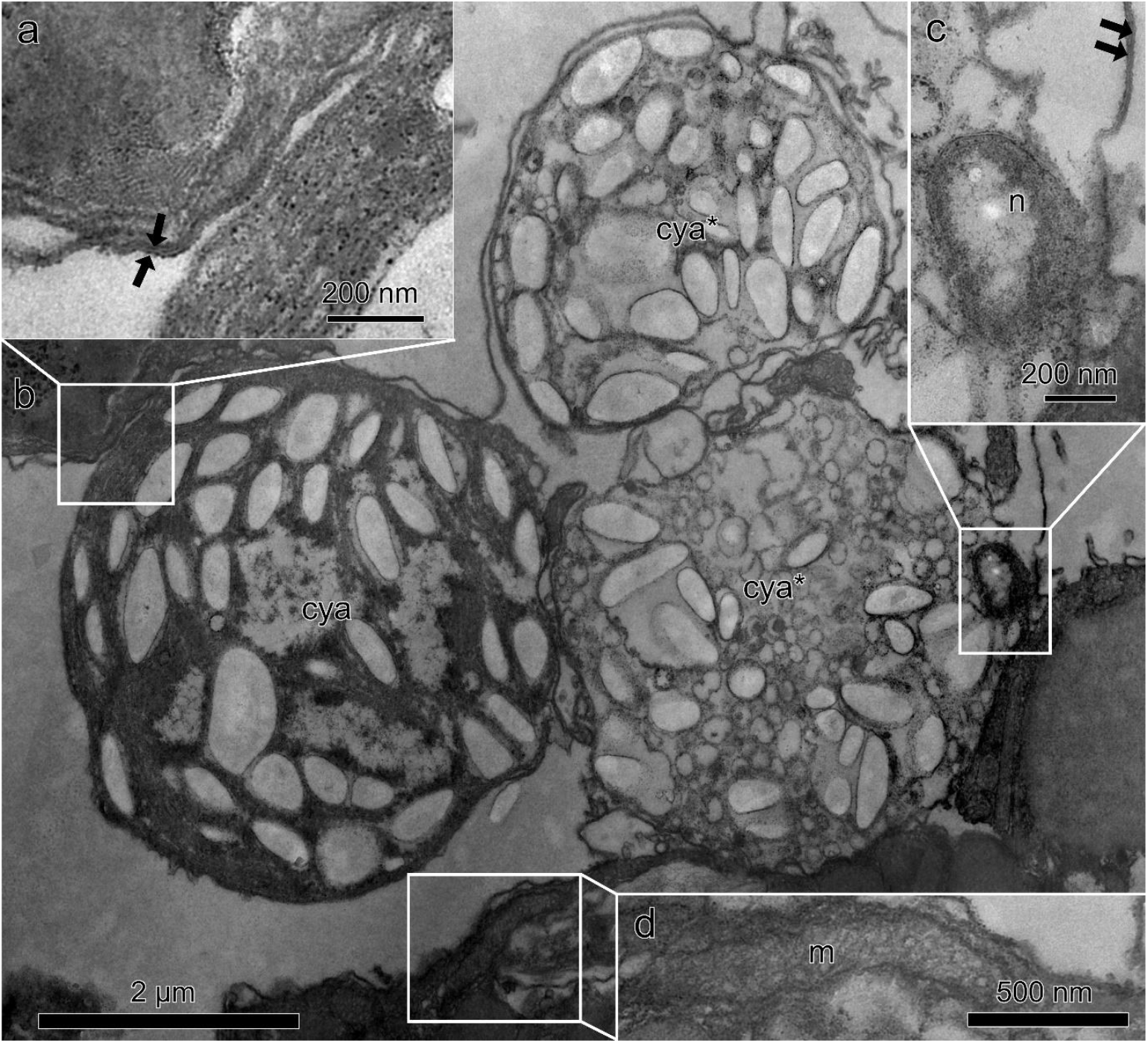
High magnification TEM section of the symbiosome in *Sinophysis* sp. **a)** Double membrane surrounding (black arrow) **b)** normal (cya) and degrading cyanobacteria (cya*), **c)** *Nisaea* (n) **d)** Mitochondria (m) on the outside.

**Figure S10.**
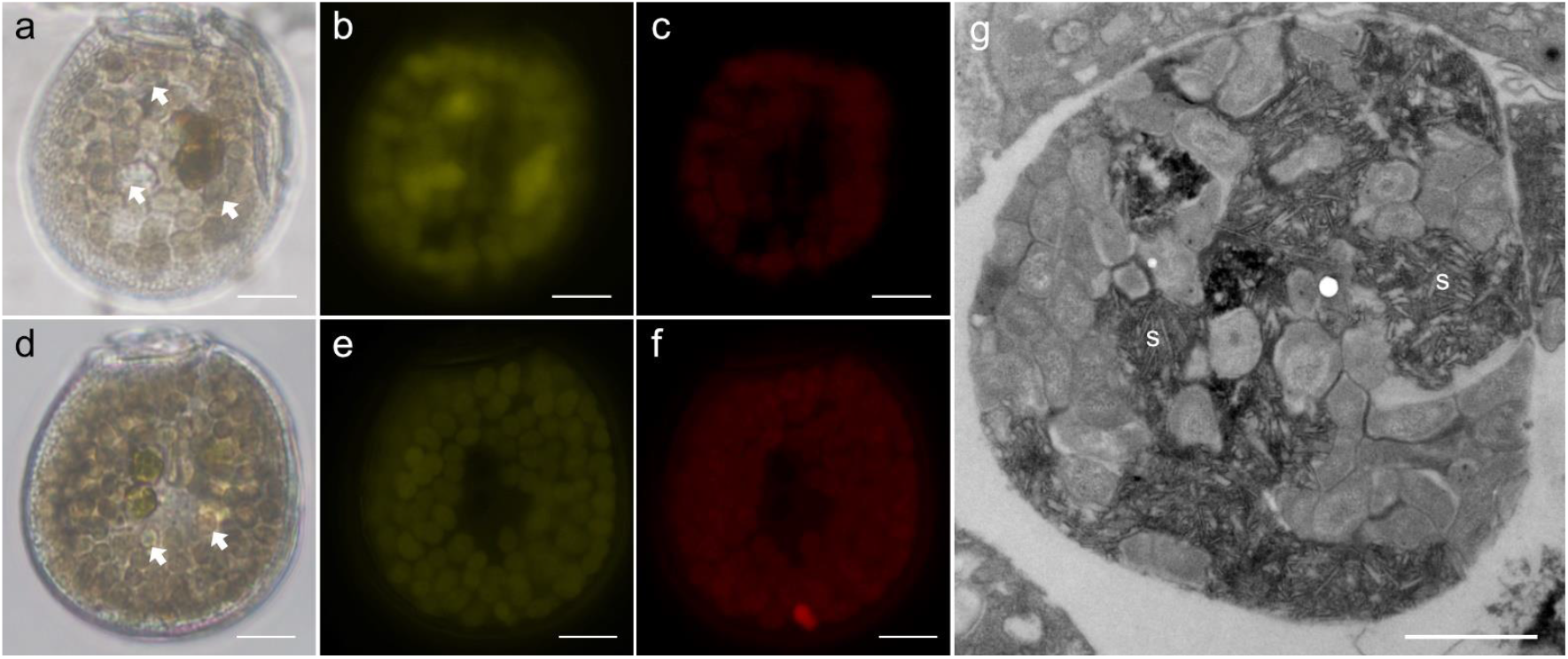
Micrographs of *Sinophysis* sp. showing the presence of food vacuoles. **a, b, c)** Lysotracker-stained cell. **d, e, f)** negative control. Food vacuole (white arrows), **g)** High magnification TEM image of a food vacuole containing minute crystalline structures (s). Scale bar = a-f) 10 µm g) 1 µm

**Figure S11.**
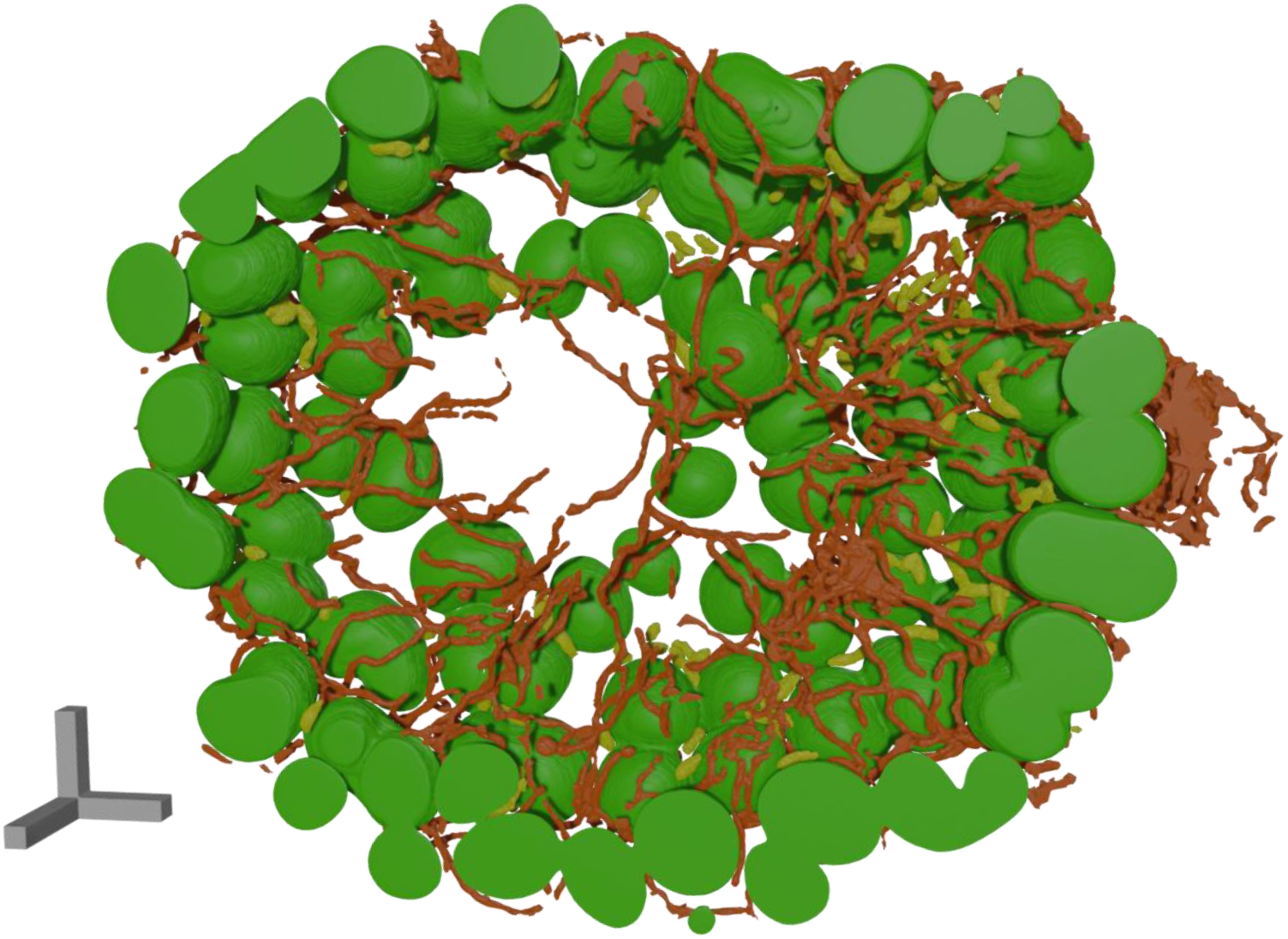
Three-dimensional reconstruction of *Sinophysis* sp. using FIB-SEM showing two symbionts. Cyanobacteria (green), alphaproteobacteria (yellow), and mitochondria (orange). Scale = 5 μm

**Figure S12.**
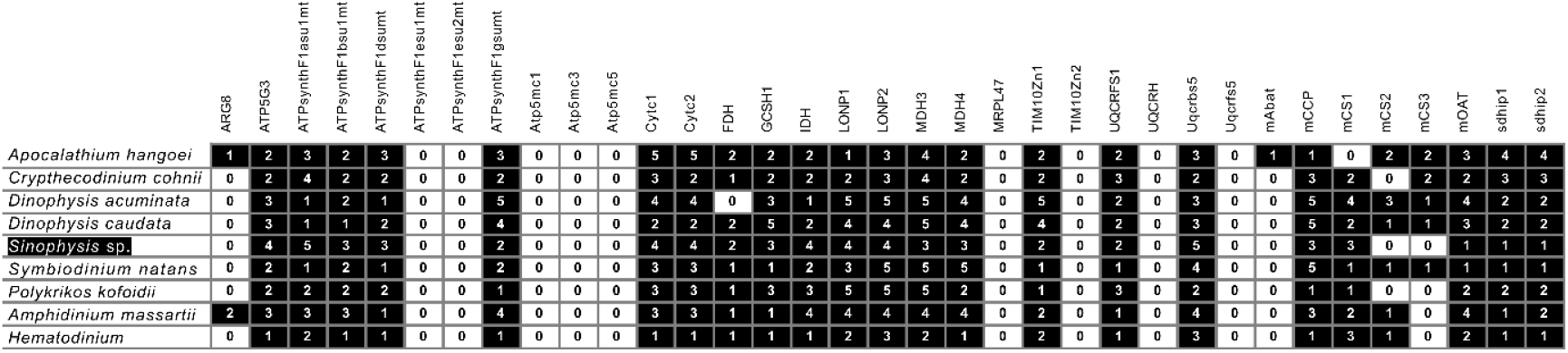
Analysis of mitochondrion-targeted proteins in *Sinophysis* sp. Oki group1. The black box indicates proteins that were identified.

**Figure S13.**
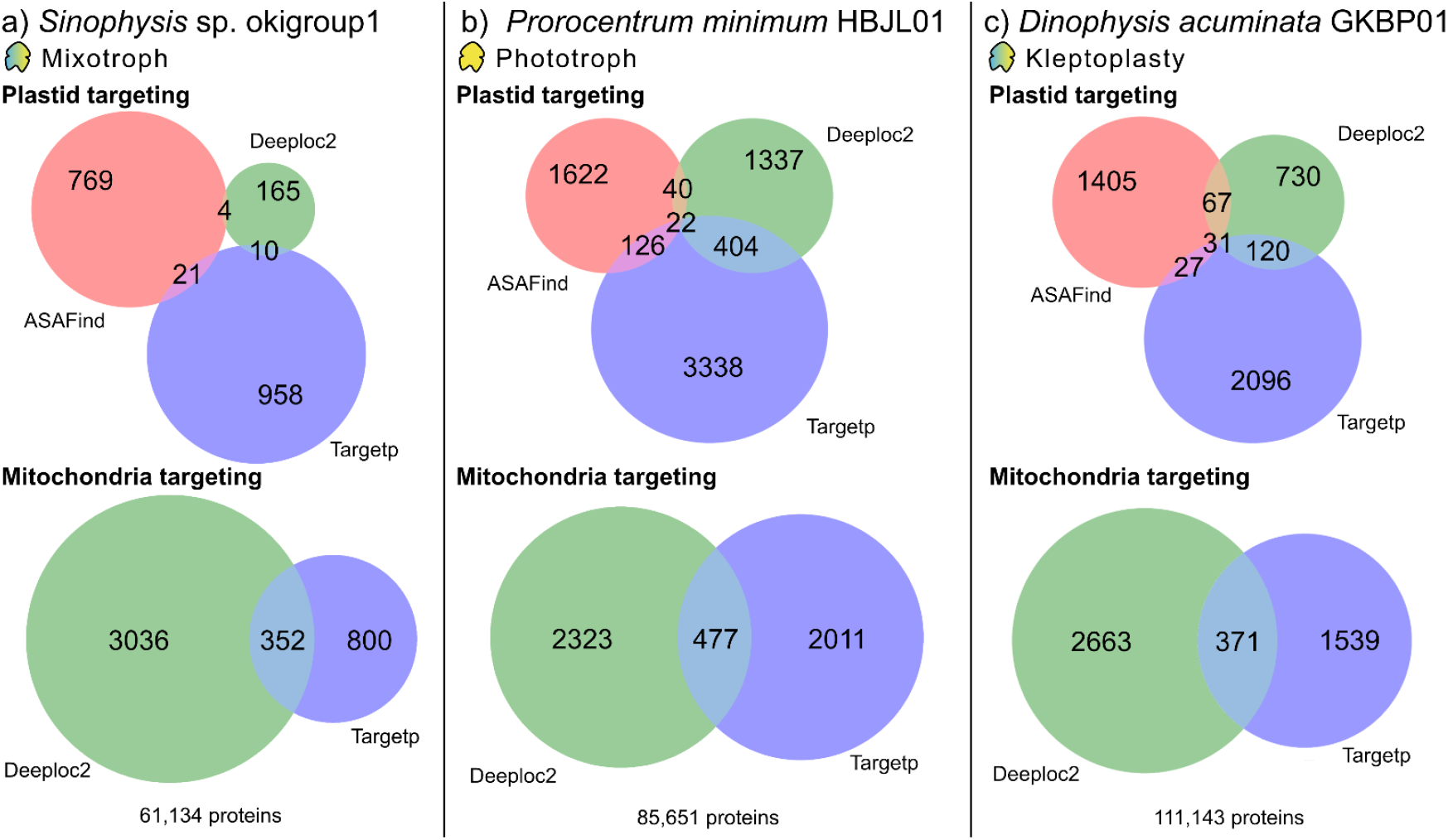
Prediction of plastid and mitochondrion targeting signal peptides from transcriptomes. **a)** *Sinophysis* sp. Okigroup1, **b)** *Prorocentrum minimum* HBJL01 and **c)** *Dinophysis acuminata* GKBP01

**Figure S14.**
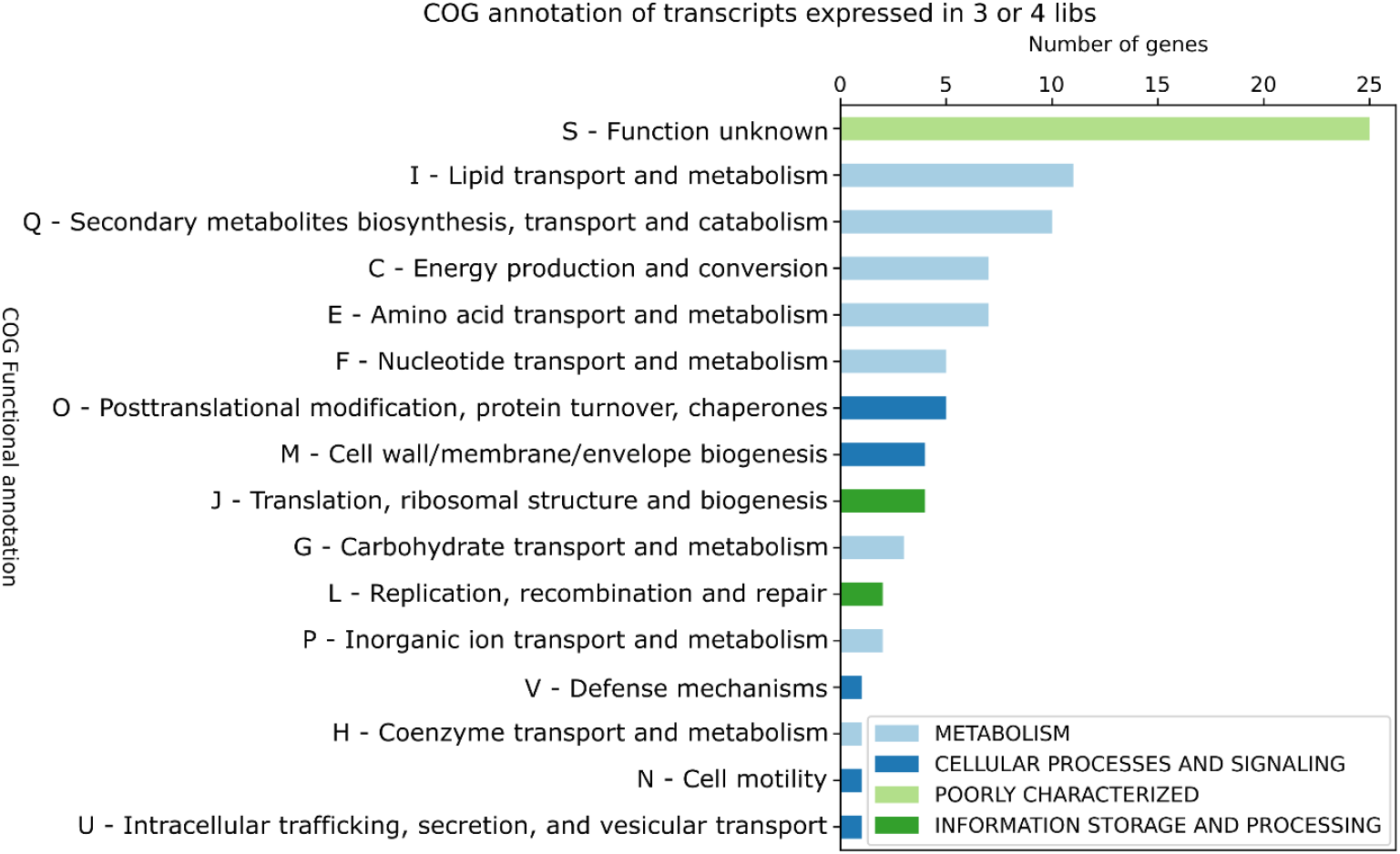
Functional COG annotation of high-confidence horizontal gene transfer candidates.

**Figure S15.**
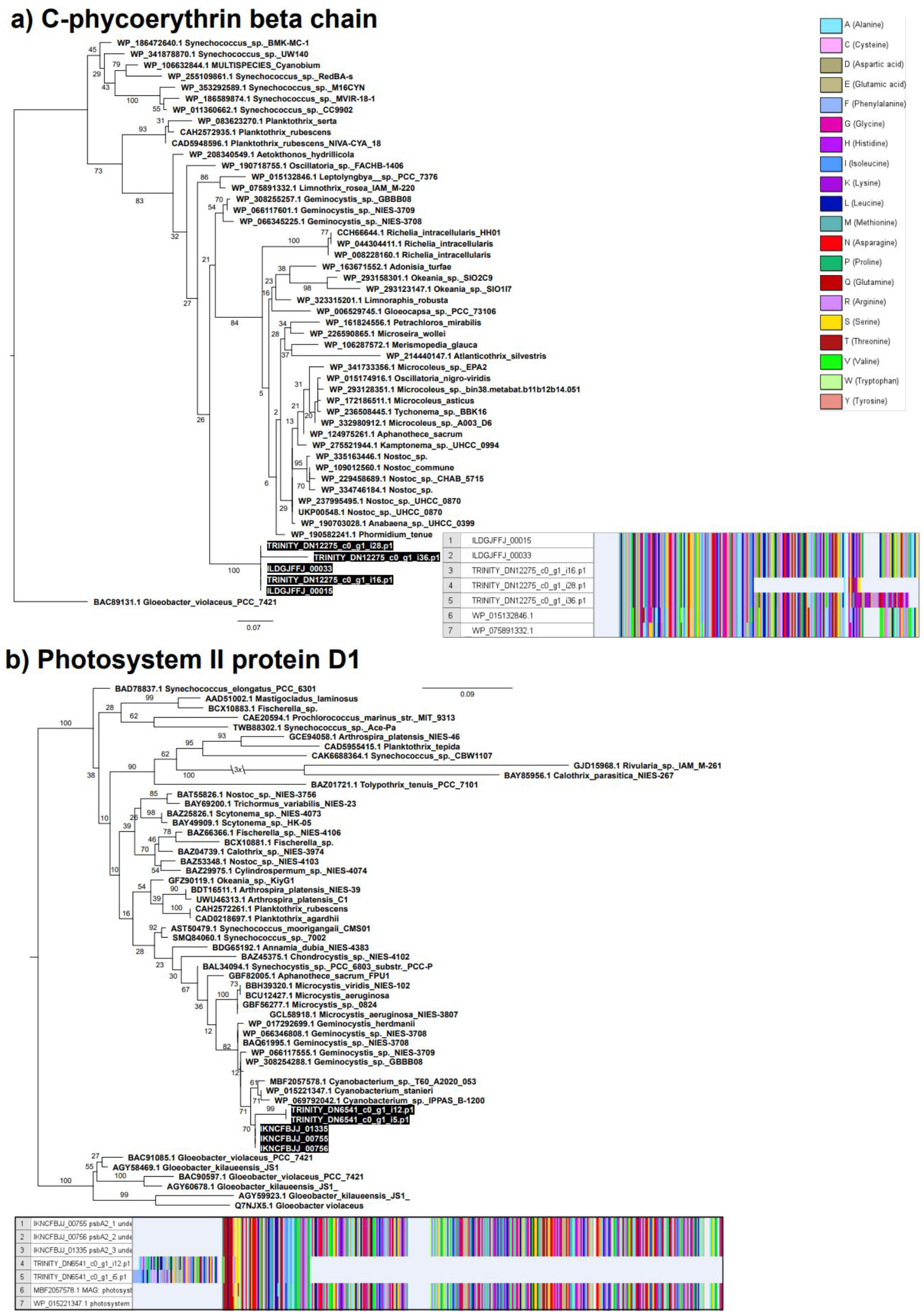
Maximum likelihood phylogenetic tree of endosymbiotic gene transfer candidates and corresponding alignments. **a)** C-phycoerythrin beta chain inferred under the Q.insect+R3 substitution model, using 1,000 bootstrap pseudo replicates. **b)** Photosystem II protein D1 inferred under the LG+R3 substitution model, using 1,000 bootstrap pseudo-replicates.

**Figure S16.**
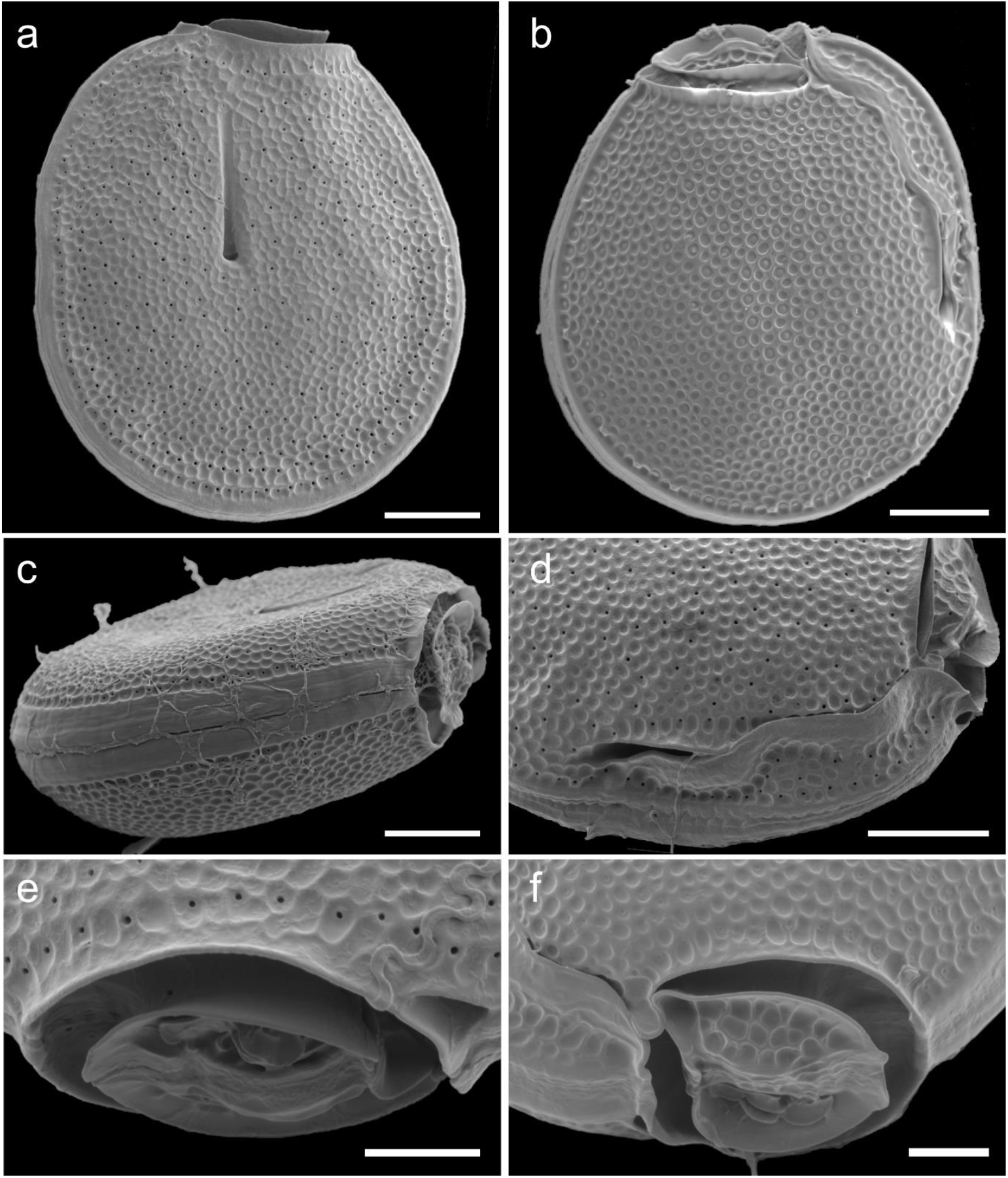
Scanning electron micrographs of *Sinophysis* sp. Okigroup1. **a)** Left lateral view, **b)** right lateral view, **c)** dorsal view, **d)** ventral view, **e)** apical view showing the flat and protruding projections, **f)** apical view showing the overlap of epitheca and hypotheca. Scale bar = a-d) 10 µm e, f) 5 µm

**Table S1.**
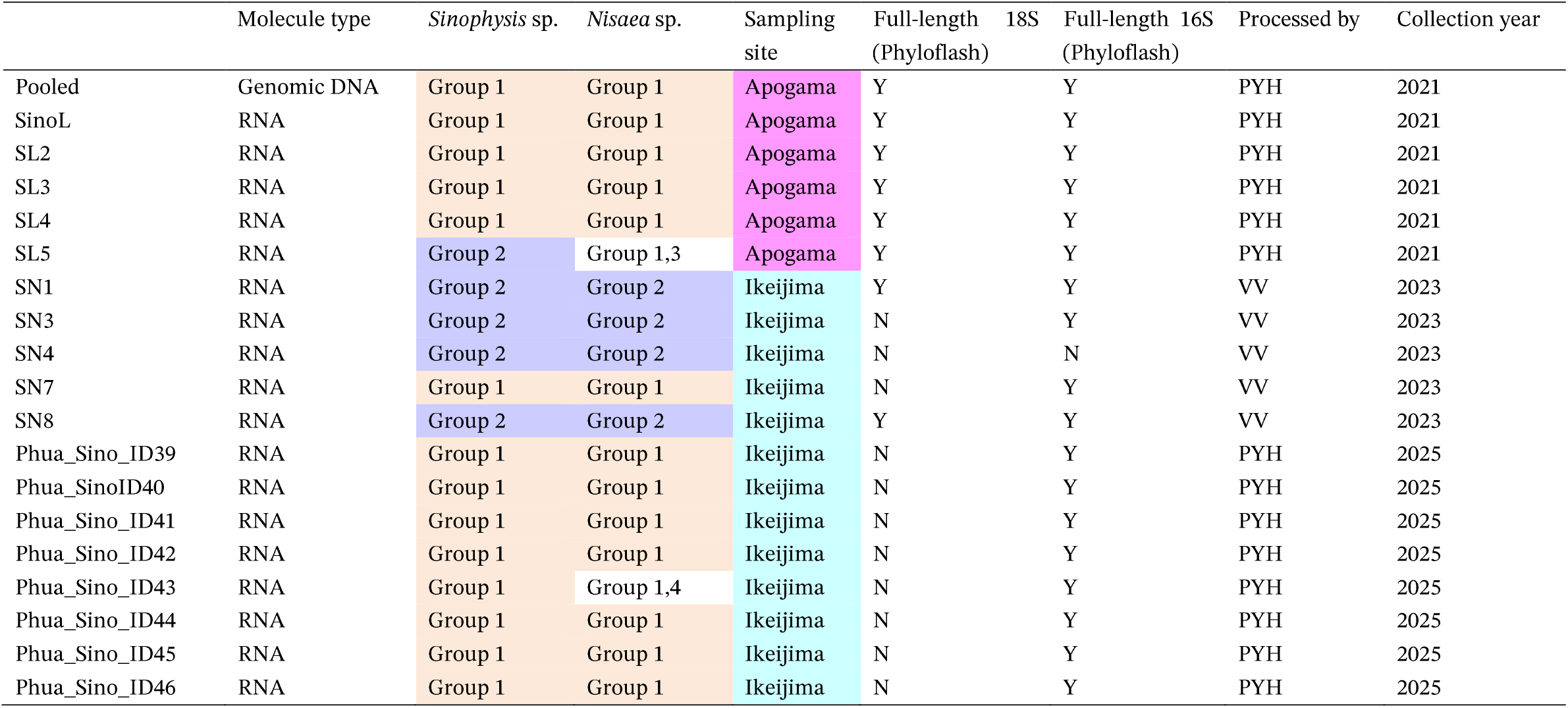
Small ribosomal subunit rDNA sequences of *Nisaea* spp. obtained from genomic (pooled sample) and single-cell libraries. For samples without full-length 16S rRNA sequences assembled via PhyloFlash, contigs from genome/transcriptome assemblies were used to extract the sequences.

**Table S2.** Overview of *Sinophysis* sp. symbiont genome features when compared to their closest free-living relatives.

| Criteria | Cyanobacterial symbiont | Alphaproteobacterial symbiont | *Geminocystis herdmanii* PCC 6308 | *Nisaea acidiphila* |
| --- | --- | --- | --- | --- |
| Genome size (bp) | 2,178,118 | 4,435,508 | 4,263,418 | 4,746,001 |
| GC content (%) | 34.7 | 64.0 | 34.3 | 63.0 |
| Number of CDS | 1678 | 3,613 | 3,981 | 4,384 |
| Number of pseudogenes | 199 | 996 | 387 | 124 |
| Number of tRNA | 42 | 39 | 40 | 51 |

**Table S5.**
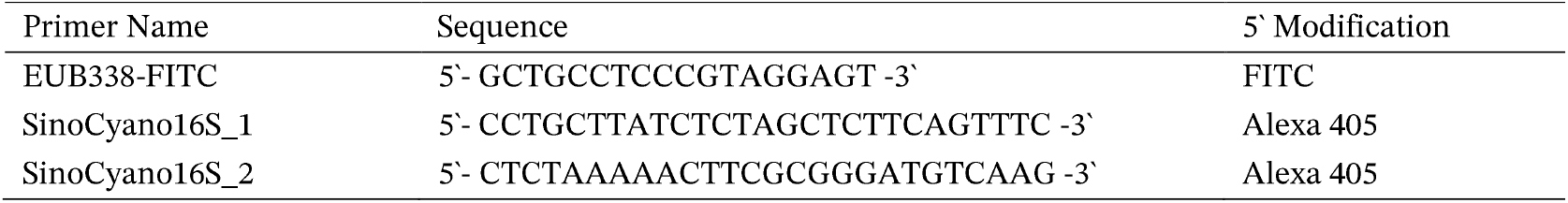
Oligonucleotide probes used for fluorescence *in situ* hybridization.

## Notes

https://github.com/ECBSU/nanorespirometry

https://github.com/ECBSU/Phua_Sinophysis_2025.git

### Competing Interest Statement

The authors have declared no competing interest.

